# Dissecting the impact of metabolic environment on three common cancer cell phenotypes

**DOI:** 10.1101/2020.06.23.167437

**Authors:** Karl Kochanowski, Timur Sander, Hannes Link, Jeremy Chang, Steven Altschuler, Lani Wu

## Abstract

The impact of different metabolic environments on cancer cell behavior is poorly understood. Here, we systematically altered nutrient composition of cell culture media and examined the impact on three phenotypes—drug-treatment survival, cell migration, and lactate overflow—that are frequently studied in cancer cells. These perturbations across diverse metabolic environments revealed simple relationships between cell growth rate and drug-treatment survival or migration. In contrast, lactate overflow was highly sensitive to changes in sugar availability but largely insensitive to changes in amino acid availability, regardless of the growth rate. Further investigation suggested that the degree of lactate overflow across metabolic environments is largely determined by the cells’ ability to maintain high rates of sugar uptake. This study enabled us to elucidate quantitative relationships between metabolic environment and cancer cell phenotypes, which echo empirical growth laws discovered to govern analogous phenotypes in microbes.

## Introduction

Cancer cells live in complex metabolic environments, encompassing a diverse range of nutrients and concentrations ^1–4^. However, little is known about the impact of metabolic environments on cancer cell behaviors. A common approach to investigate the impact of the metabolic environment *in vitro* is to subject cancer cells to pairs of defined culture media in which the concentration of one individual component has been altered. Such efforts have for example been instrumental in elucidating how changes in methionine ^5,6^, glucose ^7^, and glutamine ^8,9^ availability impact cancer cell growth. However, the extent to which changes in metabolic environment affect phenotypes other than growth is often unclear. Moreover, a limited number of pairwise comparisons may not provide insight into how complex metabolic environments affect cancer cell behaviors.

Recent advances in microbiology identified overarching relationships between metabolic environment and microbial behavior by examining cells across diverse nutrient conditions ^10–14^. Inspired by these advances, here we established an experimental workflow to produce over 100 unique *in vitro* metabolic environments by systematically altering nutrient composition. Using this workflow, we examine the impact of metabolic environment on three cancer cell phenotypes commonly studied in the context of cancer metabolism: survival of drug treatment ^15–18^, cell migration ^19,20^, and lactate overflow ^21,22^. Our data reveal simple quantitative relationships between metabolic environments and cell phenotype, which in some cases—but not all—correlate with growth rate. By exploring exceptions to these relationships, we show that the propensity for lactate secretion is primarily determined by whether the environment enables high sugar uptake rates rather than the growth rate. Overall, this work provides a framework to disentangle the complex interplay between metabolic environment, growth, and cancer cell behavior.

## Results

### Experimental workflow to generate defined metabolic environments

To enable systematic examination of the relationship between *in vitro* metabolic environment and cancer cell behavior, we adapted a synthetic media formulation ^23^ and established a two-pronged experimental workflow (Figure 1A, see methods). First, we used automated liquid transfer to generate arrays of culture media with distinct nutrient composition in 96- or 384-well plate format, which allowed cells to be exposed to multiple metabolic environments in parallel. Second, we used time-lapse microscopy to monitor growth and viability of cell populations in these media.

**Figure 1.**
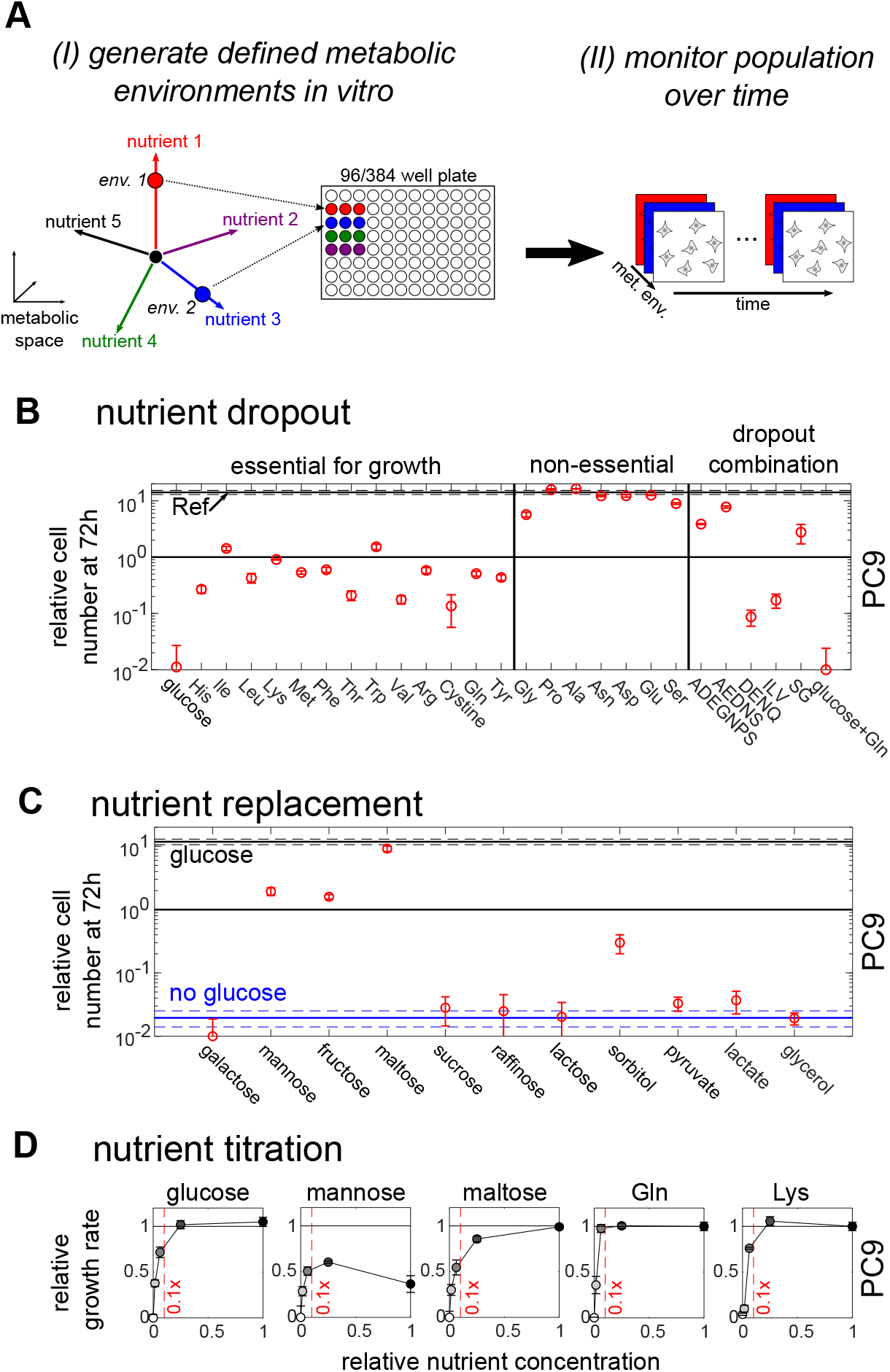
Experimental platform to systematically generate defined metabolic environments *in vitro*. **A)** Schematic of approach. **B)** Impact of dropping out individual nutrients, as well as nutrient combinations. Data shown: number of live cells at 72h relative to initial cell number. Ref: reference condition (all nutrients present at concentration corresponding to standard RPMI 1640 media). Error bars denote standard deviation (n = 2). Dropout combinations of amino acids are denoted by their single-letter code. **C)** Impact of replacing one nutrient for another, using glucose as an example. Data shown: number of live cells at 72h relative to initial cell number. Blue: relative cell number in absence of glucose. Black: relative cell number in presence of glucose (= Reference condition). Red: relative cell number in absence of glucose, but presence of the noted alternative nutrient. Error bars denote standard deviation (n = 3). **D)** Impact of gradually titrating a single nutrient on the growth rate in PC9 cells. Data shown: Monod plots of growth rate (relative to reference condition) as a function of initial nutrient concentration. Red vertical lines: concentration at 10% of reference condition as a visual aid. Error bars denote standard deviation (n = 3).

To benchmark this workflow, we used two adherent human cancer cell lines with different tissues of origin: PC9 (Non-small-cell lung cancer) and A375 (Melanoma), established *in vitro* models of EGFR- and BRAF-driven cancer, respectively ^24–26^. Specifically, we examined the impact of omitting (Figure 1B, Figures S1 and S2), replacing (Figure 1C, Figures S3 and S4), or titrating (Figure 1D, Figures S5 and S6) individual nutrients or nutrient combinations in the growth media, using a standard tissue culture medium (RPMI 1640) as a reference point. Altogether, our workflow enabled measurement of cell growth and viability for over 100 distinct nutrient compositions (list of used nutrients in supplementary tables 1 and 2).

The data recapitulated many qualitative findings from previous reports, such as the well-established essentiality of 14 nutrients for mammalian cell growth *in vitro* ^27^ (glucose and 13 amino acids; Figure 1B, Figures S1 and S2) and the ability of nutrients commonly present in the diet or cancer environment ^23^ to support growth (galactose, mannose, fructose, maltose) or extend cell survival without growth (sorbitol) when glucose is absent (Figure 1C, Figures S3 and S4). In addition, these data enabled us to observe relationships between metabolic environments and cell growth that are difficult to detect without systematic, high-throughput metabolic perturbations. First, the impact of nutrient dropout on cell survival varied considerably among the 14 essential nutrients (typically within 10-fold range, with some more extreme deviations), as well as between the two cell lines (e.g. glucose and threonine in Figure S1). Second, for most essential nutrients the maximal growth rate only diminished once nutrient concentrations were reduced more than 10-fold compared to standard cell culture conditions (Figure 1D, Figures S5 and S6), highlighting the importance of examining nutrients over a wide range of concentrations. These data demonstrate our ability to systematically and quantitatively examine cellular responses across multiple metabolic environments *in vitro* and show the wide range of cell growth rates supported by these diverse conditions.

### Impact of metabolic environment on three common cancer phenotypes

To what extent do these different metabolic environments affect phenotypes other than cell growth? We chose to investigate three phenotypes that are frequently studied in the context of cancer metabolism and can be readily quantified *in vitro*: survival of cancer drug treatment ^15–18^, cell migration ^19,20^, and lactate overflow ^21,22^ (i.e. the rapid conversion of glucose to lactate in presence of oxygen ^28^). For all three phenotypes, both metabolism and cell growth have been implicated as determinants of cell behavior. For example, slow growth has been associated with increased tolerance to both chemotherapeutics ^29,30^ and targeted therapy ^31,32^, and lactate overflow is considered a common feature of many fast-growing cell types ^28,33^. However, the extent and direction of these growth-phenotype associations are still largely unclear. For example, both negative ^34–36^ and positive ^37^ associations between growth rate and cell migration have been reported in literature.

To explore the relationship between metabolic environment, growth, and cell phenotype in a constant genetic background, we quantified each phenotype in one cell line (PC9) across ~20 diverse metabolic environments that support a wide range of growth rates. Each phenotype was quantified using recently established experimental approaches ^21,38,39^. Specifically, we examined: **i**) survival of cancer drug treatment ^16,18^ by quantifying the fraction of dead cells in a population (termed “lethal fraction”) over time with fluorescent microscopy ^38^ (Figure 2A, see methods); **ii**) cell migration by automatically tracking 39 the undirected movement of hundreds of individual cells with high-temporal resolution microscopy (Figure 3A, see methods); and **iii**) lactate overflow by quantifying lactate secretion rates using a recently published experimental protocol ^21^ (Figure 4A, see methods).

**Figure 2.**
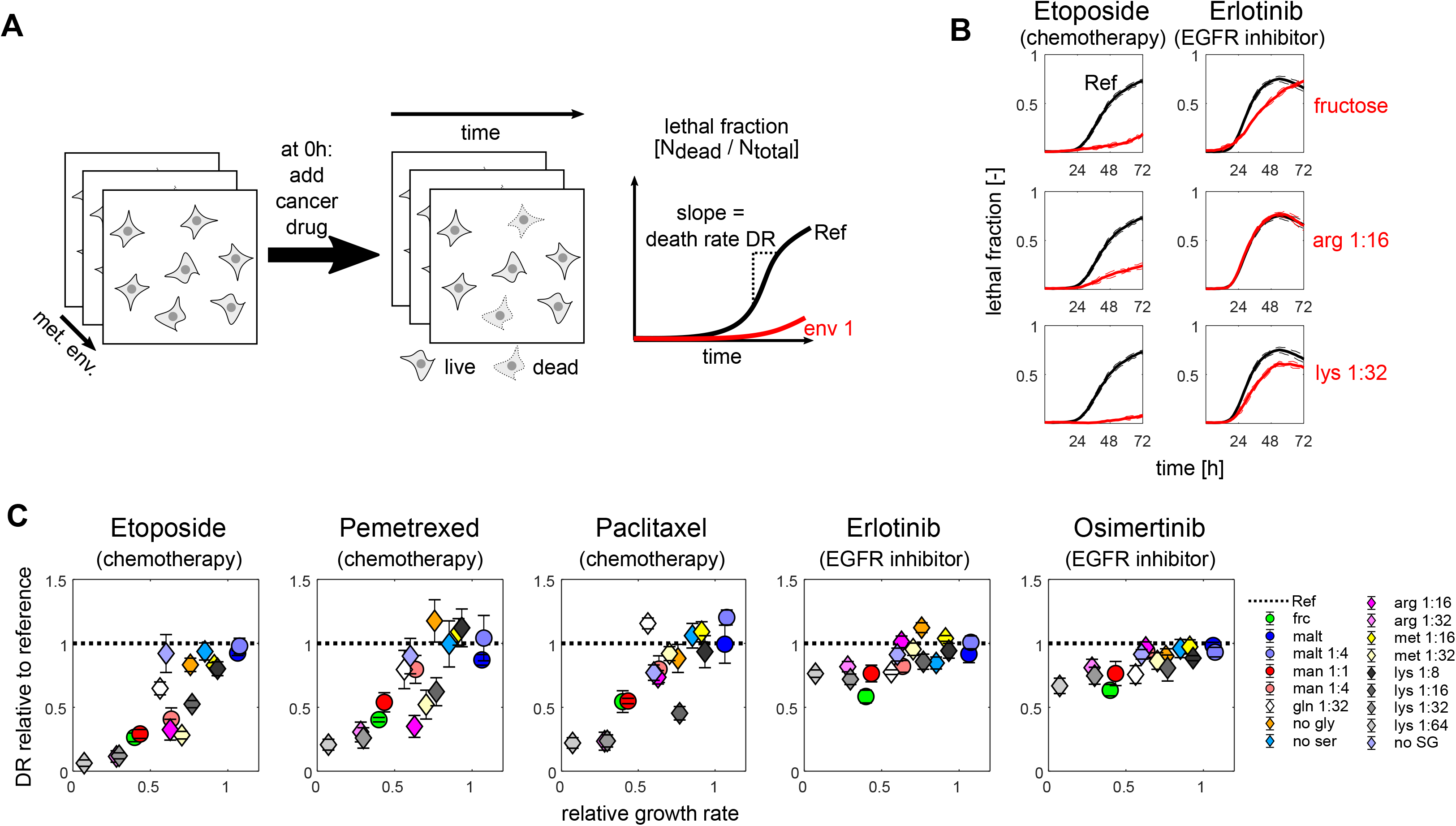
Impact of metabolic environment on cancer drug survival in PC9 cells. **A)** Schematic of experimental approach to quantify efficacy of cancer drug treatment across different metabolic environments. **B)** Time courses of PC9 lethal fraction in three different slow-growing metabolic environments (fructose, low arg/lys concentration) in presence of either Etoposide (left) or Erlotinib (right). Black: reference condition (all nutrients present at concentration corresponding to standard RPMI 1640 media). Red: respective metabolic environment. Dashed lines denote standard deviation (n = 3). **C)** Death rate (relative to reference condition) for five cancer drugs across different metabolic environments plotted as a function of growth rate (relative to reference condition). Error bars denote standard deviation (n = 3).

**Figure 3.**
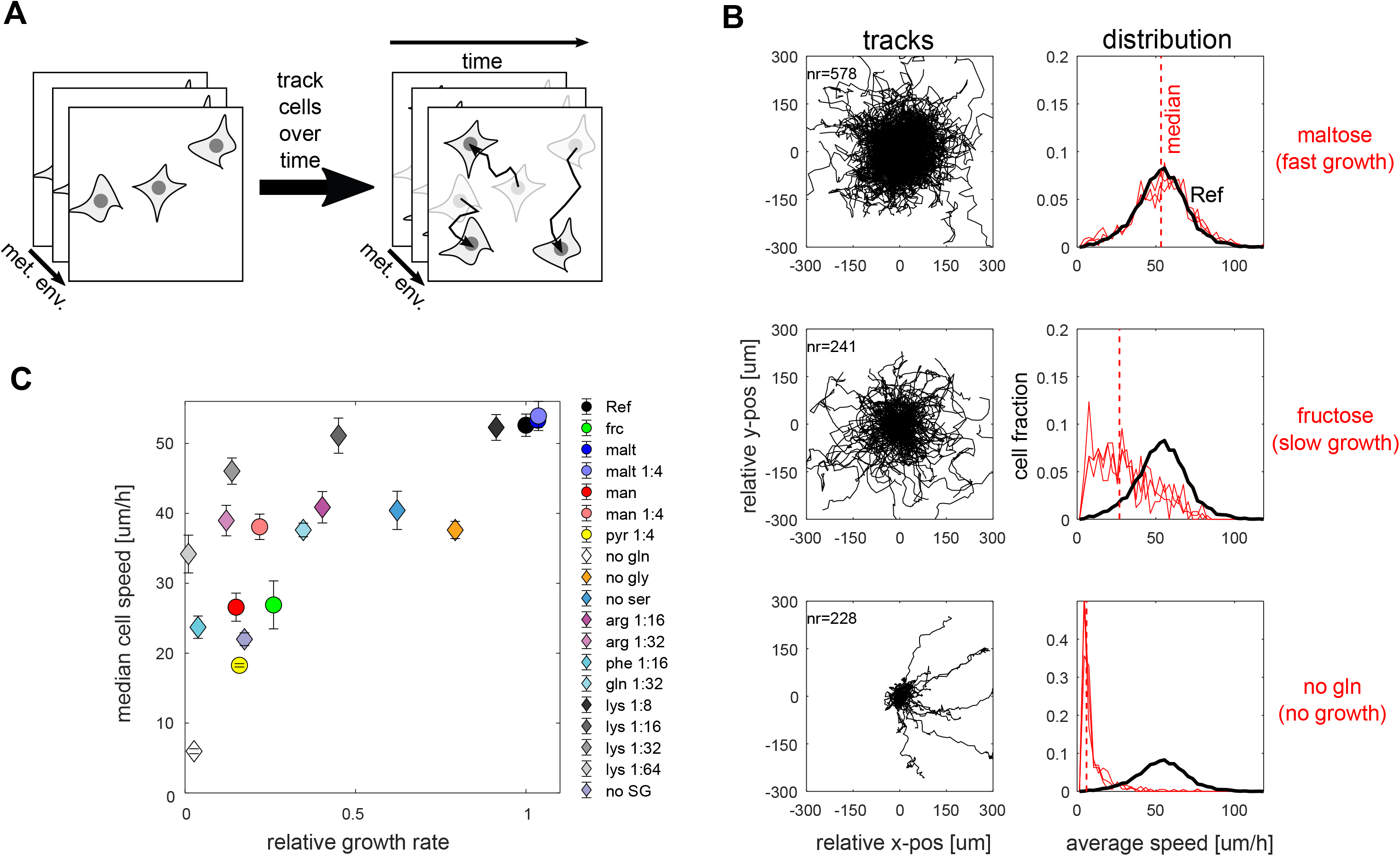
Impact of metabolic environment on cancer cell migration in PC9 cells. **A)** Schematic of experimental approach to quantify cell migration across different metabolic environments based on individual cell tracks. **B)** Left row: rose plots individual tracks captured (with track number n) over 18h in three example environments. Each track was realigned to the origin. Right row: corresponding distribution of average speed across tracks. Black: reference condition (all nutrients present at concentration corresponding to standard RPMI 1640 media). Red: designated metabolic environments. Shown are distributions of three separate replicates. Red dashed line: median of distribution across all replicates. **C)** Population speed (median of the average speed shown in B) plotted against corresponding growth rate in 19 metabolic environments. Error bars denote standard deviation (n = 3).

**Figure 4.**
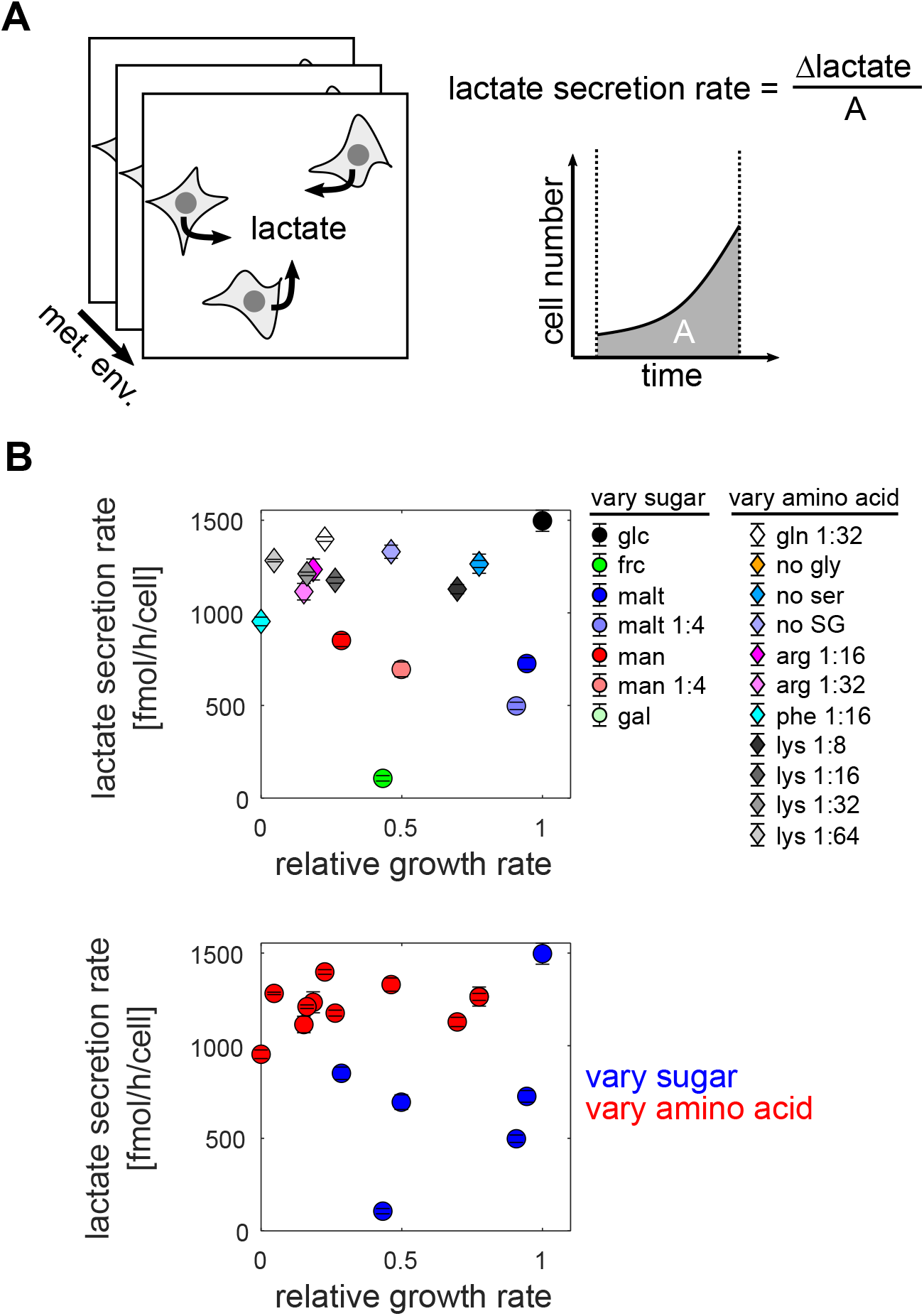
Impact of metabolic environment on lactate overflow metabolism in PC9 cells. **A)** Schematic of experimental approach to quantify lactate secretion rate. **B)** Lactate secretion rate as a function of growth rate relative to reference condition (all nutrients present at concentration corresponding to standard RPMI 1640 media) across metabolic environments in PC9 (left) and A375 (right) cells. Top row: each condition labeled separately. Bottom row: same data, but labeling conditions according to whether they constitute variation in amino acid (red) and sugar (blue) composition. Error bars denote standard deviation (n = 3).

First, we examined the impact of the metabolic environment on the efficacy of four cancer therapeutics with distinct modes of action. With our experimental approach, we measured cell survival in high doses of three commonly used chemotherapeutics (topoisomerase inhibitor Etoposide, folate antimetabolite Pemetrexed, microtubule inhibitor Paclitaxel) and the EGFR inhibitor Erlotinib as an example of targeted therapy ^26^. In metabolic environments that resemble standard tissue culture conditions, all tested drugs readily killed most cells within 72h (maximal lethal fraction approx. 0.5 or higher; black lines in Figure S7). However, the impact of the metabolic environment varied substantially across the selected drugs (examples in Figure 2B, red lines in Figure S7). In all three tested chemotherapeutics, there were dramatic differences in the rate of cell death (see methods) across conditions. As expected from previous reports comparing the chemotherapeutic sensitivity of tumors growing at different rates ^29^, the rate of cell death in chemotherapy was strongly linked with the growth rate of untreated cells in the respective metabolic environment (Figure 2C). Other metrics of drug efficacy, such as the drug-induced-proliferation rate ^40^, yielded comparable results (Figure S8). However, the rate of cell death during Erlotinib treatment was only moderately altered across conditions. We corroborated this result with the third-generation EGFR inhibitor Osimertinib, which has a distinct mode of action (covalent binding to EGFR; Figure 2C), suggesting that the metabolic environment has a moderate impact on EGFR inhibitor efficacy. Together, these data suggested that the well-established negative relationship between growth rate and cancer drug sensitivity across genetic backgrounds ^29^ holds true in cells of a common genetic background grown across multiple different metabolic environments. However, the extent of this effect depends on the cancer drug being used, with chemotherapeutic efficacy being more sensitive to metabolic environment than targeted EGFR inhibition.

Second, we examined the impact of metabolic environment on cell migration. As reported previously ^41^, PC9 cells were highly motile in metabolic environments that resemble standard tissue culture conditions (black line in right panel of Figure 3B and Figure S9). However, there was substantial variability in cell motility across metabolic environments (Figure 3B, Figure S9). The most extreme change in cell motility was observed upon complete glutamine dropout (10-fold reduction in the median cell speed of the population, Figure 3B, bottom row), echoing previous reports showing that glutaminase inhibition reduces breast cancer cell migration ^42^. Moreover, the median cell speed across metabolic environments showed significant positive correlation with the respective rate of cell growth during the experiment (Spearman correlation 0.74, p-value 4.5E-4, Figure 3C). This pattern was robust to changes in the number of cells seeded, suggesting that it is not merely a function of cell density (Figure S10). Thus, these data suggest that the metabolic environment does not affect cell migration through a “grow-**or**-go” trade-off ^34^. Instead, these data are consistent with a “grow-**and**-go” scheme, in line with recent work reporting a positive correlation between growth rate and cell migration across a panel of cancer cell lines in a fixed environment ^37^.

Third, we examined the impact of the metabolic environment on lactate overflow (Figure 4A). As with the cancer drug survival and cell migration phenotypes, we found substantial variability in lactate secretion rate across metabolic environments (Figure 4B). However, this variability was less linked to the growth rate, and instead depended on the type of nutrient varied. Environments with altered amino acid concentration showed high rates of lactate secretion regardless of growth rate (red circles in Figure 4B bottom row). In contrast, environments in which glucose had been replaced with alternative sugars showed a wide range of lactate secretion rates (blue circles in Figure 4B bottom row). Thus, our data suggest that lactate overflow in cancer cells is mostly sensitive to changes in sugar availability.

Overall, by quantifying the behavior of the same cell line across multiple different metabolic environments, we were able to identify patterns relating *in vitro* metabolic environment and cancer cell phenotypes. For drug survival and cell migration, the impact of metabolic environment on phenotype was linked to the growth rate supported by the respective environment. In contrast, lactate overflow was strongly affected by changes in sugar availability, but insensitive to changes in amino acid availability irrespective of growth rate.

### Follow-up on lactate overflow: assessing generality of patterns and identifying underlying mechanism

In the remainder of this study, we aimed to examine the relationship between metabolic environment and lactate overflow in more detail. To test whether the observed lactate overflow pattern across metabolic environments is specific to PC9 cells, we quantified lactate secretion rates in the same metabolic environments for three additional cancer cell lines with different tissues of origin and driver mutations (A375, A549, SKBR3) and one non-cancerous cell line (HEK293T). In all cell lines, the same basic pattern was conserved: amino acid variation had limited impact on lactate secretion rate regardless of supported growth rate, whereas there were wide ranges in lactate secretion depending on the supplied sugar (Figure 5). Moreover, we observed matching changes in the uptake rate of the respective sugars (Figure S11), resulting in a strong proportional relationship between lactate secretion and sugar uptake (Figure S12). That is, in metabolic environments with low lactate secretion cells also have low sugar uptake. These findings extend recent reports showing that lactate secretion and glucose uptake correlate across genetically diverse cell lines grown in a fixed metabolic environments ^8,21,22^. Our data show that this positive correlation is also maintained across sugars in a fixed genetic background, and reaffirm the notion that lactate is largely derived from glycolysis ^28^.

**Figure 5.**
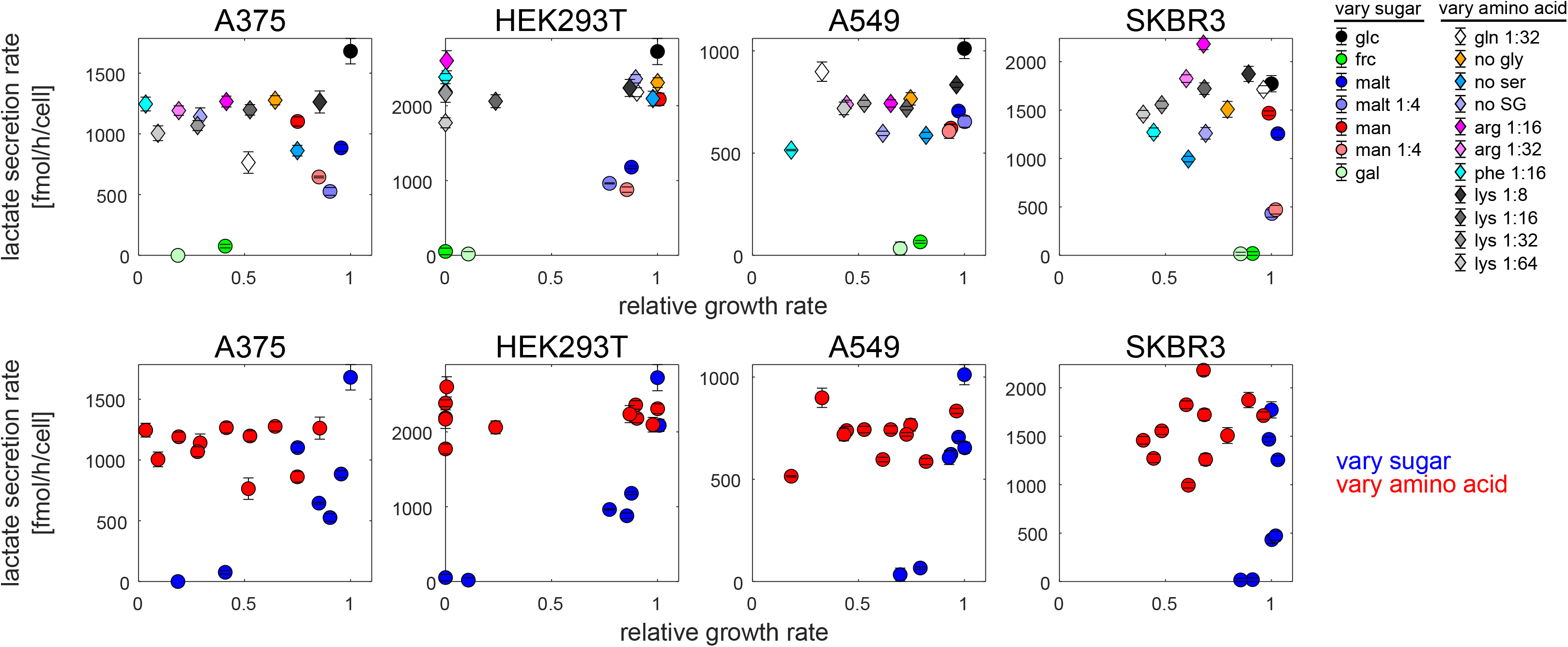
Lactate overflow pattern across metabolic environments is conserved in diverse cell lines. Lactate secretion rate as a function of growth rate relative to reference condition (all nutrients present at concentration corresponding to standard RPMI 1640 media) across metabolic environments in four additional cell lines. Top row: each condition labeled separately. Bottom row: same data, but labeling conditions according to whether they constitute variation in amino acid (red) and sugar (blue) composition. Error bars denote standard deviation (n = 3).

A possible explanation for the low lactate secretion rates in metabolic environments such as fructose and galactose is that in these cases ATP production through respiration (which has a higher ATP yield than glycolysis, Figure 6A) already matches the cell’s ATP demand. As a result, additional ATP production through increased glycolysis would be unnecessary. To test this hypothesis, we quantified the ratio of AMP to ATP (a measure of the energetic state of the cell) in metabolic environments with low/intermediate/high rates of lactate secretion in two cell lines (PC9, A375) using targeted metabolomics. Overall, both cell lines showed a very similar metabolome response in the different metabolic environments (supplementary Figure S13). Importantly, the AMP to ATP ratio showed a negative relationship with the lactate secretion rate across metabolic environments (Figure 6B), as did e.g. the GMP to GTP ratio (supplementary Figure S14). These results suggested that cells growing in environments with low lactate secretion rates are, in fact, energy limited.

**Figure 6.**
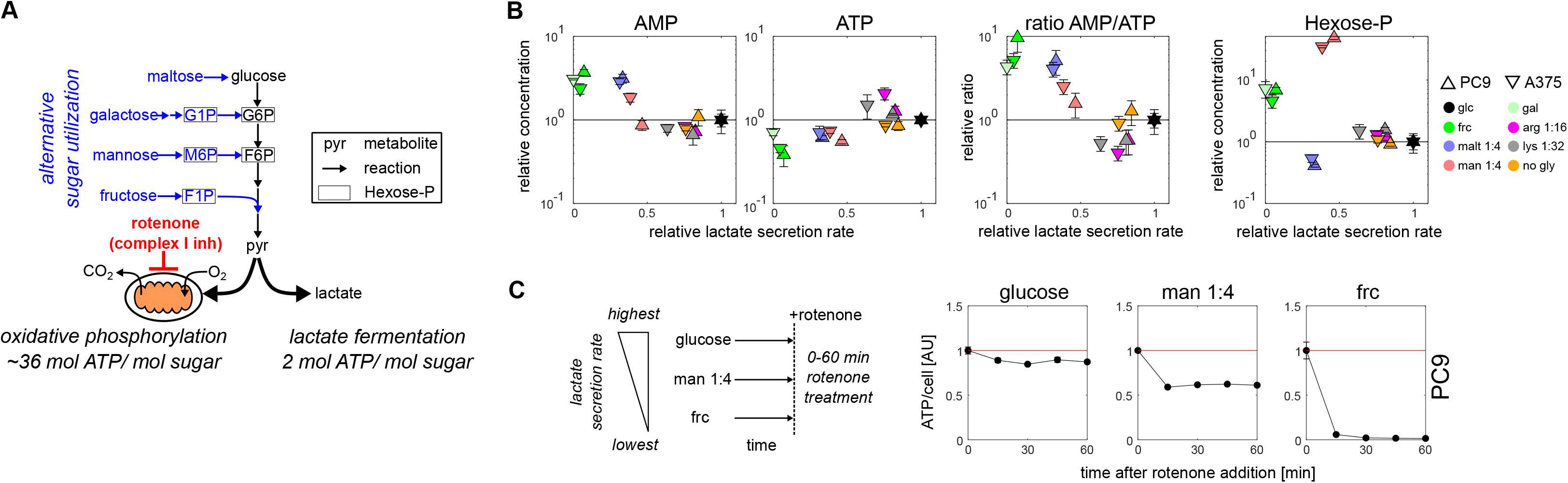
Investigating the source of variability in lactate secretion rate across metabolic environments. **A)** Schematic of mammalian sugar utilization and catabolism by oxidative phosphorylation or lactate fermentation. Boxed metabolites (e.g. G6P) denote hexose-phosphates. **B)** Intracellular concentration of (left to right) AMP, ATP, ratio of AMP/ATP relative to reference condition, and hexose-P plotted against the corresponding lactate secretion rate (relative to reference condition) in PC9 and A375 cells. Error bars denote standard deviation (n = 3). **C)** ATP concentration (determined with CellTiterGlo, normalized to signal at t = 0 min) upon complex I inhibition with 1 uM rotenone in PC9 cells growing in conditions with high (glucose), intermediate (1:4 man), or low (frc) lactate secretion rates. Error bars denote standard deviation (n = 2).

Why do cells growing in these metabolic environments with alternative sugars such as fructose not simply increase glycolysis to meet their energy demands? These alternative sugars have distinct entry routes into glycolysis (Figure 6A), which rely on metabolic reactions (i.e. transport- and internal conversion reactions) unique to each sugar. A parsimonious explanation is that metabolic bottlenecks somewhere along these unique entry routes render cells unable to increase glycolysis to match their energy demands. While pinpointing metabolic bottlenecks remains challenging even in microbial systems ^43^, we reasoned that intracellular metabolite data may enable us to distinguish between two potential bottlenecks for each sugar: the internal conversion step (which often manifests as an accumulation of the immediate substrate) or the sugar transport (where no such accumulation is observed). Our metabolomics data cannot distinguish the hexose-phosphate species unique to each sugar utilization pathway (Figure 6A). Nevertheless, the dramatic increase in the total hexose-phosphate pool during growth on galactose, mannose and fructose (Figure 6B) pointed towards internal bottlenecks, as recently suggested for galactose-grown HEK293T cells ^44^. In contrast, the lack of hexose-phosphate accumulation for maltose rather pointed towards transport bottlenecks.

To test the hypothesis that cells are unable to increase the glycolysis of alternative sugars such as fructose, we examined the impact of electron transfer chain inhibition, which forces cells to increase glycolysis to maintain energy balance. During growth on glucose, cells compensated for rotenone inhibition of complex I inhibition by increasing the rate of sugar uptake (Figure S15), allowing cells to maintain high ATP levels (Figure 6C, left). In contrast, rotenone treatment in metabolic environments with intermediate/low lactate rates (mannose and fructose) caused a dramatic drop in ATP (Figure 6C, middle and right), in line with previous reports ^45^. We note that we could mimic this drop in ATP with rotenone treatment during growth on glucose by inhibiting sugar uptake with the specific GLUT1 inhibitor Bay876 (Figure S16), which by itself does not affect ATP levels (Figure S17). Taken together, our results suggest that the low lactate secretion rates observed in slow-growth metabolic environments such as fructose or galactose are driven by the cells’ inability to increase glycolysis due to metabolic bottlenecks in the respective utilization pathways.

## Discussion

Here, we examined the impact of metabolic environment on three common cancer cell phenotypes: drug survival, cell migration, and lactate overflow. Quantifying cell behavior in different metabolic environments—with varying sugar and amino acid concentrations—revealed simple relationships that often correlated with growth rate. For example, in metabolic environments that only supported slow growth, cells showed higher resilience to chemotherapeutics and reduced motility. In contrast, lactate overflow was sensitive to changes in available sugars, but largely insensitive to changes in amino acid concentration regardless of the growth rate.

The phenotype relationships across metabolic environments for the tested cancer cells reported here are remarkably similar to the empirical “growth laws” ^46,47^ by which nutrient availability governs analogous phenotypes in microbes. For example, slow-growth environments tend to increase antibiotic survival ^48,49^ and decrease colony expansion rates ^50^ in bacteria. This resemblance is particularly striking when comparing the patterns of overflow metabolism in cancer cells and microbes: as observed here in cancer cells, the main determinant of microbial overflow metabolism is also not growth rate *per se*, but rather the type of nutrient limitation cells face in their environment ^11,51^. Thus, our findings suggest that the phenotypic response of cancer cells to changes in metabolic environment may be shaped by similar metabolic constraints as those found in bacteria, and may adhere to similar empirical “growth laws”. Moreover, our findings have implications beyond cancer, since the many of the phenotypes investigated here are not exclusive to cancer cells. For example, lactate overflow is found in many other fast growing cell types, such as activated T-cells ^52^. While we have used cancer cells as a test case, this work may guide future efforts to examine whether the metabolic environment exerts similar effects on the behavior of other cell types.

One notable exception from these general relationships was the impact of the metabolic environment on EGFR inhibitor efficacy (see Figure 2). Despite recent reports highlighting the importance of metabolism for targeted therapy survival ^16,53,54^, EGFR inhibitor efficacy changed only moderately across metabolic environments for comparable drug concentrations. A possible explanation for this discrepancy is that the targeted metabolic perturbations typically used in literature (i.e. direct inhibition of metabolic enzymes with small-molecule compounds ^16,53,54^) may trigger changes in metabolic activity that are distinct from the changes in metabolic environment used here. Moreover, even though prior literature had hinted towards a negative correlation between cell growth and targeted therapy survival ^31,32^, the data presented here suggest that, at least in the context of EGFR inhibition, metabolically induced slow growth is not sufficient to dramatically increase EGFR inhibitor survival. Future efforts may test the extent to which these findings generalize to other EGFR inhibitors or other models of targeted therapy.

There are several limitations of this present study. First, is the selection of metabolic environments, which focused on varying amino acid and sugar concentration. Clearly, this selection is not exhaustive, and there are other nutrients present *in vivo* ^23,55^ (lipids, dicarboxylic acids, vitamins) which could be explored in future studies. Moreover, although using an established cell culture formulation (RPMI 1640) as a reference point facilitated comparison with prior literature, the nutrient concentrations cancer cells encounter *in vivo* are likely to differ substantially. Nevertheless, as quantitative *in vivo* information about the metabolic environment of tumors becomes available ^4^, this study may serve as a template for identifying changes in microenvironment that are critical for a phenotype of interest.

Second, we did not tackle the question of how different genetic backgrounds affect the observed metabolic environment-phenotype relationships. Our data suggests that the metabolic environment may affect lactate secretion rates in similar ways across diverse cell lines. However, it is not clear whether the same is true for drug survival or cell migration. Future efforts may use the approaches presented here to examine the impact of genetic background (i.e. different driver mutations) on the relationship between metabolic environment and these phenotypes, and potentially other phenotypes of interest.

Finally, while we focused on the phenotypic characterization of cells across metabolic environments, we did not assess how the relationship between metabolic environment and observed phenotype is established mechanistically. For example, we did not address the mechanisms underlying the differences in cell motility observed here (Figure 3). Previous work has highlighted the importance of ATP production for cell motility ^19,20^. Consistently, we find that inhibition of ATP production has a dramatic effect on PC9 cell motility (Figure S18). Another candidate mechanism is mTOR, which acts as a nutrient sensor ^56,57^, and whose inhibition also impairs cell motility in PC9 cells (Figure S18). For lactate overflow, our data already point towards a potential mechanism, namely that lactate secretion is determined not by growth rate, but rather by the cells’ ability to maintain high rates of sugar uptake. This would be consistent with recent work showing that glucose import is a key flux-controlling step in glycolysis ^58^. Moreover, the elevated AMP to ATP ratio (indicating energy limitation) we observed in environments with low sugar uptake suggests that the additional ATP provided by elevated glycolysis constitutes a major contribution to the cell’s energy balance.

In conclusion, our study provides a quantitative framework for systematically examining the impact of different metabolic environments on cancer cell behavior. This allowed us to uncover relationships between growth rate and different cancer cell phenotypes. The patterns we uncover provide a step towards elucidating phenotypic “growth laws” of cancer, as has been studied in bacteria, and guide future mechanistic efforts.

## Supporting information

composition of reference media

additional nutrients used in study

## Acknowledgments

The authors thank Alain Bonny for assistance with FACS sorting. K.K. received postdoctoral fellowships from the European Molecular Biology Organization (long-term fellowship ALTF 1167–2016) and the Swiss National Science Foundation; S.J.A. and K.K.’s training were partially supported by GM112690 to S.J.A.; L.F.W. is supported by NCI-NIH R01 CA184984. Thanks to Mattia Zampieri, Maike Roth, Heinz Hammerlindl, Xiaoxiao Sun, Louise Heinrich, and Leanna Morinishi for their feedback on the manuscript.

## Author contributions

Conceived the study: KK, LW, SA. Performed experiments: KK, TS, JC. Analyzed data: KK, TS, HL. Wrote manuscript: KK, LW, SA, with input from all authors.

## Methods

### Cell lines

PC9 cells were obtained from the Minna Laboratory at UT Southwestern, and all other cell lines used in here were obtained from the UCSF cell culture facility. PC9, A375, HEK293T, and SKBR3 cell lines harboring H2B-mCherry were constructed using lentiviral gene transfer. H2B-mCherry transfer vector was a gift from the Yang lab at Columbia University. After transfection, cells harboring mCherry were FACS sorted. Cell identity was confirmed using STR profiling, and cells were examined and found negative for mycoplasma contamination.

### Media

Cell lines were maintained in phenol-red free RPMI 1640 media (Gibco, 11835-030) supplemented with 5% fetal bovine serum (Gemini Bioproducts, CAT 100-106, LOT A15G00I) and 1% antibiotic-antimycotic (Gemini Bioproducts, CAT 400-101, LOT F23S00J), which was sterile-filtered (0.22 μm membrane, Olympus, CAT 25-227) before usage.

Unless stated otherwise, all experiments were performed in defined synthetic cell culture media, termed reconstituted media, based on a recently published formulation ^23^, with the following modifications: First, amino acids were used at the same concentration as in standard RPMI 1640 media with exception of alanine (not included in RPMI 1640 media, used at 0.1 mM) and arginine (used at 0.115 mM). Second, additional polar metabolites listed in the published formulation ^23^ were omitted. Third, glucose was used at 5.55 mM concentration (half the concentration of RPMI 1640 media). Fourth, phenol red was omitted to avoid interference with fluorescence imaging. Finally, Sytox Green (Life Technologies, CAT S7020, used at 20 nM) was added to detect dead cells based on their green fluorescent signal. See table S1 for the exact media composition. In experiments involving nutrient replacement, the respective nutrient was omitted and replaced with one alternative nutrient per condition. See table S2 for the full list of alternative nutrients. All media and nutrients were sterile-filtered (0.22 μm membrane, Olympus, CAT 25-227) before usage.

### Cultivation

Cell cultivation was performed as follows: Cultures grown in maintenance media as described above (to 60-80% confluence) were trypsinized for 5 min at 37C (0.25% Trypsin, Gemini Bioproducts, CAT 400-151), centrifuged (RT, 300 g, 3 min), and re-suspended in pre-heated reconstituted media lacking amino acids and glucose.

For benchmarking and drug survival experiments, nutrients were transferred to 384-well plates (Corning, CAT 353962) using an ECHO liquid handler (Labcyte ECHO 525/650) to the designated concentrations, and cell suspensions were added (50 μL culture volume, seeding 1000 cells per well). Plates were sealed with BreathEasy foil (Neta Scientific, CAT RPI-248738) to minimize evaporation and incubated at 37C with 5% CO_2_ for 16-20h before starting the time course experiments to allow cells to adhere to the plate bottom. In the case of drug survival experiments, cancer drugs (or DMSO as solvent control) were added 1:1000 using an ECHO liquid handler immediately before starting the time lapse microscopy experiments.

For cell migration experiments, nutrients were transferred to 96-well IncuCyte ImageLock plates (Essen Biosciences, CAT 4379) using an ECHO liquid handler (Labcyte ECHO 525/650) to the designated concentrations, and cell suspensions were added (100 μL culture volume, seeding 3000 cells per well). Plates were sealed with BreathEasy foil (Neta Scientific, CAT RPI-248738) to minimize evaporation and incubated at 37C with 5% CO_2_ for 16-20h before starting the time lapse microscopy experiments to allow cells to adhere to the plate bottom.

For lactate overflow, metabolomics, and acute rotenone treatment experiments, nutrients were transferred to 96-well plates (Corning, CAT 353219) using an ECHO liquid handler (Labcyte ECHO 525/650) to the designated concentrations, and cell suspensions were added (100 μL culture volume, seeding 5000 cells per well). Plates were sealed with BreathEasy foil (Neta Scientific, CAT RPI-248738) to minimize evaporation and incubated at 37C with 5% CO_2_ for 12h before starting the time lapse microscopy experiments to allow cells to adhere to the plate bottom.

### Time lapse microscopy

Cell cultures were monitored over time (at 37C and 5% CO_2_) using the IncuCyte S3 automated imaging system (Essen Biosciences): at regular intervals (every 2h for benchmarking, drug survival, and lactate overflow experiments, every 15 min for cell migration experiments) phase/RFP/GFP images were taken of each well (one image per well, settings: 10x objective, 300/400 ms acquisition time for GFP/RFP). From these images, the total number of cells (defined as the number of objects in the RFP channel) and the number of dead cells (defined as the number of objects which overlap in RFP and GFP channels) were extracted using built-in IncuCyte Analysis Software. The number of live cells per image and time point was calculated as the total number of cells minus the number of dead cells, and normalized to the first time point to yield time courses of relative cell numbers. The lethal fraction per image and time point was calculated as the number of dead cells divided by the total number of cells^38^. For A549 cells (which did not harbor H2B-mCherry), confluence in phase images was converted to cell number using a separate calibration curve obtained with rapid-red nuclear dye (Essen Biosciences, CAT 4706, used 1:2000).

### Quantification of growth and death rates

Growth rates was calculated from cell number time courses following previously published approaches for cancer ^40^ and microbial ^14,59^ cell cultures. Briefly, time-dependent growth rate μ(t) was estimated by linear regression of the relative cell number measurement (in log scale) within a sliding window of 9 consecutive time points (corresponding to a 16h time window, equivalent to the approximate doubling time of PC9 and A375 cells in standard media). The maximal value of μ(t) across the time course was defined as the maximal growth rate μ_max_ of the respective culture. Conversely, the time-dependent death rate DR(t) was estimated by linear regression of the lethal fraction measurement within a sliding window of 9 consecutive time points. The maximal value of DR(t) across the time course was defined as the death rate of the respective culture. All calculations were performed using MatLab (MatLab 2019a).

### Quantification of cell motility from time course experiments

Motility of individual cells from time lapse microscopy experiments was quantified based on a previously published single-particle tracking algorithm^39^ using custom-made MatLab scripts (MatLab 2019a). First, stacks of RFP images (18h duration, imaging interval 15 min) were imported for each well, and Nuclei were identified by LoG detection (MatLab command *edge*, LoG threshold 0.025). Starting from the first time point, x/y positions of nuclei centroids were extracted in consecutive pairs of images (MatLab command *regionprops*). Next, the Euclidean distance between each centroid in image 1 and image 2 was calculated (Matlab command *pdist2*) and used to solve the linear assignment problem (MatLab command *matchpairs*, cost of being unmatched: 20). Single cell tracks were constructed from links between centroids in consecutive images, only considering tracks with a length > 10 consecutive time points (= 2.5h). The average speed of each track was calculated as the mean Euclidean distance across all time points within a track, and converted into the final unit (micrometer/h) based on the objective-specific conversion factor of 1.24 μm/pixel and a frame rate of 1 image every 15 min. To obtain a population level metric of cell motility, the median of average track speeds within the well was calculated.

### Quantification of lactate secretion and sugar uptake

Lactate secretion and sugar uptake rates were quantified following a previously published approach^21^. Briefly, lactate secretion was quantified as follows: First, the amount of lactate, which had been produced over the course of the experiment (36-48h experiment duration), was quantified from culture supernatants using a commercial lactate quantification kit (Megazyme, CAT K-LATE) in 96-well plate format following the manufacturers’ protocol. Second, the lactate secretion rate was calculated by normalizing the molar amount of lactate produced by the area under the growth curve^21^ (calculated from cell number time courses quantified as described above).

Similarly, sugar uptake was determined by first quantifying the amount of sugar, which had been consumed over the course of the experiment (i.e. initial sugar concentration minus remaining sugar concentration at sampling time), from culture supernatants using commercial enzymatic assays (Megazyme, CAT K-GLUC for glucose; CAT K-MANGL for mannose/fructose; CAT K-ARGA for galactose; CAT K-MASUG for maltose) in 96-well plate format following the manufacturers’ protocol. The sugar uptake rate was then calculated by normalizing the molar amount of sugar consumed by the area under the growth curve, as described for the lactate secretion rate quantification.

### Metabolomics

Intracellular metabolite concentrations was quantified following previously published protocols ^22,60^. Briefly, cells were cultivated in 96-well plate format as described above in the designated metabolic environments for at least 24h. Subsequently, media were removed, cells were washed once with 100 μL 75 mM ammonium carbonate (pH 7.4, 37 °C), and 100 μL quenching/extraction solution was added (40% methanol, 40% acetonitrile, 20% water, −20 °C). Plates were kept at −20 °C for 2h, centrifuged (4000 rpm, 5 min), and 50 μL metabolite extracts were transferred to conical storage plates (ThermoFisher, CAT AB-1058), sealed (ThermoFisher, CAT AB-0745) and kept at −80 °C. Prior to measurement, 50 μL metabolite extracts of E. coli grown on ^13^C glucose were added to serve as internal ^13^C standard^60^. Metabolite concentrations in extracts were then quantified by liquid chromatography coupled to tandem mass spectrometry (LC−MS/MS) as described before ^60^, and for each metabolite and sample peak intensity was normalized to the respective 13C peak intensity and the confluence (at time of extraction as a proxy for biomass).

### Response to acute rotenone treatment

Cells were cultivated in 96-well plate format as described above in the designated metabolic environments for at least 24h. Rotenone was then added to a final concentration of 1 μM. After 15 to 60 mins, ATP levels (proxy for the cellular energetic state) were quantified with CellTiter-Glo 2.0 (Promega, CAT G9243) following the manufacturers’ protocol. ATP signals were normalized to DMSO-treated controls grown in the respective metabolic environments.

## References

1. Hensley, C. T. et al. Metabolic Heterogeneity in Human Lung Tumors. Cell 164, 681–94 (2016).

2. Kamphorst, J. J. et al. Human pancreatic cancer tumors are nutrient poor and tumor cells actively scavenge extracellular protein. Cancer Res. 75, 544–53 (2015).

3. Reznik, E. et al. A Landscape of Metabolic Variation across Tumor Types. Cell Syst. 1–13 (2018). doi:10.1016/j.cels.2017.12.014

4. Sullivan, M. R. et al. Quantification of microenvironmental metabolites in murine cancers reveals determinants of tumor nutrient availability. Elife 8, 1–27 (2019).

5. Mentch, S. J. et al. Histone Methylation Dynamics and Gene Regulation Occur through the Sensing of One-Carbon Metabolism. Cell Metab. 22, 861–873 (2015).

6. Wang, Z. et al. Methionine is a metabolic dependency of tumor-initiating cells. Nat. Med. 25, (2019).

7. Birsoy, K. et al. Metabolic determinants of cancer cell sensitivity to glucose limitation and biguanides. Nature 508, 108–12 (2014).

8. Chen, P.-H. et al. Metabolic Diversity in Human Non-Small Cell Lung Cancer Cells. Mol. Cell 76, 838–851.e5 (2019).

9. Timmerman, L. A. et al. Glutamine Sensitivity Analysis Identifies the xCT Antiporter as a Common Triple-Negative Breast Tumor Therapeutic Target. Cancer Cell 24, 450–465 (2013).

10. You, C. et al. Coordination of bacterial proteome with metabolism by cyclic AMP signalling. Nature 500, 301–306 (2013).

11. Basan, M. et al. Overflow metabolism in Escherichia coli results from efficient proteome allocation. Nature 528, 99–104 (2015).

12. Nichols, R. J. et al. Phenotypic landscape of a bacterial cell. Cell 144, 143–56 (2011).

13. Schmidt, A. et al. The quantitative and condition-dependent Escherichia coli proteome. Nat. Biotechnol. 34, 104–10 (2016).

14. Kochanowski, K. et al. Few regulatory metabolites coordinate expression of central metabolic genes in Escherichia coli. Mol. Syst. Biol. 13, 903 (2017).

15. Viswanathan, V. S. et al. Dependency of a therapy-resistant state of cancer cells on a lipid peroxidase pathway. Nature 547, 453–457 (2017).

16. Momcilovic, M. et al. Targeted Inhibition of EGFR and Glutaminase Induces Metabolic Crisis in EGFR Mutant Lung Cancer. Cell Rep. 18, 601–610 (2017).

17. Viale, A. & Draetta, G. F. Metabolic features of cancer treatment resistance. in Recent Results in Cancer Research 207, 135–156 (2016).

18. Hardeman, K. N. et al. Dependence on Glycolysis Sensitizes BRAF-mutated Melanomas for Increased Response to Targeted BRAF Inhibition. Sci. Rep. 7, 1–9 (2017).

19. Zanotelli, M. R. et al. Regulation of ATP utilization during metastatic cell migration by collagen architecture. Mol. Biol. Cell 29, 1–9 (2018).

20. Yizhak, K. et al. A computational study of the Warburg effect identifies metabolic targets inhibiting cancer migration. Mol. Syst. Biol. 10, 744 (2014).

21. Jain, M. et al. Metabolite Profiling Identifies a Key Role for Glycine in Rapid Cancer Cell Proliferation. Science (80-.). 336, 1040–1044 (2012).

22. Ortmayr, K., Dubuis, S. & Zampieri, M. Metabolic profiling of cancer cells reveals genome-wide crosstalk between transcriptional regulators and metabolism. Nat. Commun. 10, 1841 (2019).

23. Cantor, J. R. et al. Physiologic Medium Rewires Cellular Metabolism and Reveals Uric Acid as an Endogenous Inhibitor of UMP Synthase. Cell 169, 258–272.e17 (2017).

24. Sharma, S. V. et al. A Chromatin-Mediated Reversible Drug-Tolerant State in Cancer Cell Subpopulations. Cell 141, 69–80 (2010).

25. Shaffer, S. M. et al. Rare cell variability and drug-induced reprogramming as a mode of cancer drug resistance. Nature 546, 431–435 (2017).

26. Kochanowski, K., Morinishi, L., Altschuler, S. & Wu, L. Drug persistence - from antibiotics to cancer therapies. Curr. Opin. Syst. Biol. 10, 1–8 (2018).

27. Eagle, H. Nutrition needs of mammalian cells in tissue culture. Science 122, 501–14 (1955).

28. Deberardinis, R. J. & Chandel, N. S. We need to talk about the Warburg effect. Nat. Metab. 2, 127–129 (2020).

29. Baguley, B. C. et al. Resistance mechanisms determining the in vitro sensitivity to paclitaxel of tumour cells cultured from patients with ovarian cancer. Eur. J. Cancer 31, 230–237 (1995).

30. Pearl Mizrahi, S., Gefen, O., Simon, I. & Balaban, N. Q. Persistence to anti-cancer treatments in the stationary to proliferating transition. Cell Cycle 15, 3442–3453 (2016).

31. Roesch, A. et al. A Temporarily Distinct Subpopulation of Slow-Cycling Melanoma Cells Is Required for Continuous Tumor Growth. Cell 141, 583–594 (2010).

32. Roesch, A. et al. Overcoming intrinsic multidrug resistance in melanoma by blocking the mitochondrial respiratory chain of slow-cycling JARID1B(high) cells. Cancer Cell 23, 811–25 (2013).

33. Vander Heiden, M. G. M. G., Cantley, L. C. L. C. & Thompson, C. B. C. B. Understanding the Warburg effect: the metabolic requirements of cell proliferation. Science 324, 1029–33 (2009).

34. Giese, A. et al. Dichotomy of astrocytoma migration and proliferation. Int. J. cancer 67, 275–82 (1996).

35. Tiek, D. M. et al. Alterations in Cell Motility, Proliferation, and Metabolism in Novel Models of Acquired Temozolomide Resistant Glioblastoma. Sci. Rep. 8, 1–11 (2018).

36. Atkins, R. J. et al. Cell quiescence correlates with enhanced glioblastoma cell invasion and cytotoxic resistance. Exp. Cell Res. 374, 353–364 (2019).

37. Garay, T. et al. Cell migration or cytokinesis and proliferation? – Revisiting the “go or grow” hypothesis in cancer cells in vitro. Exp. Cell Res. 319, 3094–3103 (2013).

38. Forcina, G. C., Conlon, M., Wells, A., Cao, J. Y. & Dixon, S. J. Systematic Quantification of Population Cell Death Kinetics in Mammalian Cells. Cell Syst. 4, 600–610.e6 (2017).

39. Jaqaman, K. et al. Robust single-particle tracking in live-cell time-lapse sequences. Nat. Methods 5, 695–702 (2008).

40. Harris, L. A. et al. An unbiased metric of antiproliferative drug effect in vitro. Nat. Methods 13, 497–500 (2016).

41. Takai, E., Tsukimoto, M., Harada, H. & Kojima, S. Autocrine signaling via release of ATP and activation of P2X7 receptor influences motile activity of human lung cancer cells. Purinergic Signal. 10, 487–497 (2014).

42. Wang, J. Bin et al. Targeting mitochondrial glutaminase activity inhibits oncogenic transformation. Cancer Cell 18, 207–219 (2010).

43. Liu, Y. et al. overproducing Bacillus subtilis. Nat. Commun. (2016). doi:10.1038/ncomms11933

44. Li, S. et al. Galactose 1-phosphate accumulates to high levels in galactose-treated cells due to low GALT activity and absence of product inhibition of GALK. 1–11 (2019). doi:10.1002/jimd.12198

45. Marroquin, L. D., Hynes, J., Dykens, J. a, Jamieson, J. D. & Will, Y. Circumventing the Crabtree effect: replacing media glucose with galactose increases susceptibility of HepG2 cells to mitochondrial toxicants. Toxicol. Sci. 97, 539–47 (2007).

46. Scott, M. & Hwa, T. Bacterial growth laws and their applications. Curr. Opin. Biotechnol. 1–7 (2011). doi:10.1016/j.copbio.2011.04.014

47. Basan, M. Resource allocation and metabolism: the search for governing principles. Curr. Opin. Microbiol. 45, 77–83 (2018).

48. Brauner, A., Fridman, O., Gefen, O. & Balaban, N. Q. Distinguishing between resistance, tolerance and persistence to antibiotic treatment. Nat. Rev. Microbiol. 14, 320–330 (2016).

49. Fung, D. K. C., Chan, E. W. C., Chin, M. L. & Chan, R. C. Y. Delineation of a bacterial starvation stress response network which can mediate antibiotic tolerance development. Antimicrob. Agents Chemother. 54, 1082–93 (2010).

50. Cremer, J. et al. Chemotaxis as a navigation strategy to boost range expansion. Nature (2019). doi:10.1038/s41586-019-1733-y

51. Brauer, M. J. et al. Coordination of growth rate, cell cycle, stress response, and metabolic activity in yeast. Mol. Biol. Cell 19, 352 (2008).

52. Brand, K. et al. Cell‐cycle‐related metabolic and enzymatic events in proliferating rat thymocytes*. Eur. J. Biochem. 172, 695–702 (1988).

53. Raha, D. et al. The Cancer Stem Cell Marker Aldehyde Dehydrogenase Is Required to Maintain a Drug-Tolerant Tumor Cell Subpopulation. Cancer Res. 74, 3579–3590 (2014).

54. Hangauer, M. J. et al. Drug-tolerant persister cancer cells are vulnerable to GPX4 inhibition. Nature (2017). doi:10.1038/nature24297

55. Voorde, J. Vande et al. Improving the metabolic fidelity of cancer models with a physiological cell culture medium. Sci. Adv. 5, (2019).

56. Zoncu, R., Efeyan, A. & Sabatini, D. M. mTOR: from growth signal integration to cancer, diabetes and ageing. Nat. Rev. Mol. Cell Biol. 12, 21–35 (2011).

57. Mossmann, D., Park, S. & Hall, M. N. mTOR signalling and cellular metabolism are mutual determinants in cancer. Nat. Rev. Cancer 18, 1 (2018).

58. Tanner, L. B. et al. Four Key Steps Control Glycolytic Flux in Mammalian Cells Article Four Key Steps Control Glycolytic Flux in Mammalian Cells. Cell Syst. 1–14 (2018). doi:10.1016/j.cels.2018.06.003

59. Balaban, N. Q. et al. Definitions and guidelines for research on antibiotic persistence. Nat. Rev. Microbiol. (2019). doi:10.1038/s41579-019-0196-3

60. Guder, J. C., Schramm, T., Sander, T. & Link, H. Time-Optimized Isotope Ratio LC–MS/MS for High-Throughput Quantification of Primary Metabolites. Anal. Chem. 89, 1624–1631 (2017).

