## supplementary figures for "Dissecting the impact of metabolic environment on three common cancer cell phenotypes"

**Figure S1. Impact of dropping out individual nutrients, as well as nutrient combinations, in PC9 (top panel) and A375 (bottom panel) cells.** Ref: reference condition (all nutrients present at concentration corresponding to standard RPMI 1640 media). Error bars denote standard deviation of 2 replicate wells. Dropout combinations of amino acids are denoted by their single-letter code. Relative cell numbers at 72h below 0.01 were set to 0.01 to aid visualization.

**Figure S2. Time courses of relative cell numbers in nutrient dropout experiments, in PC9 (A) and A375 (B) cells.** Ref: reference condition (all nutrients present at concentration corresponding to standard RPMI 1640 media). Dashed lines denote standard deviation of 2 replicate wells. Dropout combinations of amino acids are denoted by their single-letter code.

**Figure S3. Impact of replacing glucose for alternative nutrients, in PC9 (top panel) and A375 (bottom panel) cells.** Blue: relative cell number at 72h in absence of glucose. Black: relative cell number in presence of glucose (= Reference condition). Red: relative cell number in absence of glucose, but presence of the noted alternative nutrient. Error bars denote standard deviation of 3 replicate wells. Relative cell numbers at 72h below 0.01 were set to 0.01 to aid visualization.

**Figure S4. Time courses of relative cell numbers in nutrient replacement experiments, in PC9 (top row) and A375 (bottom row) cells.** Blue: absence of glucose. Black: glucose (= Reference condition). Red: absence of glucose, but presence of the noted alternative nutrient. Dashed lines denote standard deviation of 3 replicate wells.

**Figure S5. Growth rate as a function of initial nutrient concentration in PC9 (A) and A375 (B) cells.** Red vertical lines: concentration at 10% of reference condition (all nutrients present at concentration corresponding to standard RPMI 1640 media) as a visual aid. Error bars denote standard deviation of three replicate wells.

**Figure S6. Time courses across nutrient titrations of PC9 (A-B) and A375 (C-D) cells.** Black: reference condition (all nutrients present at concentration corresponding to standard RPMI 1640 media). Red: metabolic environments with nutrient limitation. **A)** Amino acid titration in PC9 cells. **B)** Sugar titration in PC9 cells. **C)** Amino acid titration in A375 cells. **D)** Sugar titration in A375 cells. Dashed lines denote standard deviation of 2-3 replicates.

**Figure S7. Lethal fraction time courses of PC9 cells across metabolic environments treated with DMSO (top row), and high doses of 5 different cancer drugs.** Black: lethal fraction time course in reference condition (all nutrients present at concentration corresponding to standard RPMI 1640 media). Red: lethal fraction time course in designated metabolic environment. Dashed lines denote standard deviation (n = 3). DMSO: 0.1%. Etoposide: 5 uM. Pemetrexed: 2 uM. Paclitaxel 5 uM. Erlotinib: 1 uM. Osimertinib: 0.1 uM.

**Figure S8. Overview of diverse metrics to quantify impact of metabolic environment on PC9 survival.** Top row: 72h viability relative to viability in reference condition treated with same drug. Second row: 72h viability relative to viability in respective drug-free (i.e. DMSO treated) metabolic environment. Third row: drug-induced-proliferation (DIP) rate (defined as maximal slope of live cell time course) relative to DIP rate in reference condition. Fourth row: Row 5: maximal LF value within 72h of drug treatment. Bottom row: relative death rate, same as in main figure 2C. Error bars denote standard deviation (n = 3). All data are plotted against relative growth rate in DMSO-treated controls (maximal growth rate in respective metabolic environment relative to reference condition, see methods for details).

**Figure S9. Distribution of average PC9 cell speeds in individual tracks across metabolic environments.** Black: reference condition (all nutrients present at concentration corresponding to standard RPMI 1640 media). Red: designated metabolic environments. Shown are distributions of three separate replicates. Red dashed line: median of distribution across all replicates. Black dashed line: median of average cell speed in Reference condition.

**Figure S10. Median cell speed at different seeding numbers.** Left: PC9 median cell speed at 3000 (x-axis) versus 5000 (y-axis) cells seeded per well (96-well plate format) across metabolic environments. Right: corresponding growth rate. Error bars denote standard deviation (n = 3).

**Figure S11. Sugar uptake rate as a function of growth rate relative to reference condition (all nutrients present at concentration corresponding to standard RPMI 1640 media) across metabolic environments in five cell lines.** Top row: each condition labeled separately. Bottom row: same data, but labeling conditions according to whether they constitute variation in amino acid (red) and sugar (blue) composition. Error bars denote standard deviation (n = 3).

**Figure S12. Amount of lactate produced plotted against the corresponding amount of consumed sugar across metabolic environments in five cell lines.** Error bars denote standard deviation (n = 3). Dashed line: 2:1 relationship between produced lactate and consumed sugar. Continuous line: 1:1 relationship.

**Figure S13. Intracellular metabolome across metabolic environments. A)** log2 fold-changes of intracellular metabolite concentration (relative to glucose condition) for PC9 (left) and A375 (right) cells. Note that PC9 cells do not grow on galactose, therefore this condition was omitted for this cell line. **B)** Same data as in A), plotting intracellular concentrations (relative to glucose) for 6 overlapping conditions against each other (x-axis: PC9, y-axis: A375). Empty circles denote metabolites, which change less than two-fold in both cell lines (highlighted by box with dashed lines). Filled circles: metabolite concentration changes > two-fold in at least one of the cell lines. Selected metabolites with large fold-changes are labeled. Error bars denote standard deviation (n = 3).

**Figure S14.** Intracellular concentration ratios of selected metabolites relative to reference condition plotted against the corresponding lactate secretion rate (relative to reference condition) in PC9 and A375 cells. Error bars denote standard deviation (n = 3).

**Figure S15. Impact of 24h treatment with metabolic effectors on lactate secretion in PC9 and A375 cells.** Cells were seeded in RPMI 1640 media with 5% FBS, and after 24h media were exchanged for fresh RPMI media containing metabolic effectors at the denoted concentrations. **A)** Lactate secretion rate of PC9 (left) and A375 (right) cells treated with complex I inhibitor Rotenone (Rot, yellow symbols). As additional controls for the experimental setup, GLUT1 inhibitor Bay876, glycolysis inhibitor 2DG, and LDH inhibitor Oxamate (Oxa) were used, which all reduce lactate secretion rate. Error bars denote standard deviation (n = 3). **B)** Lactate secretion rate plotted against corresponding glucose uptake rate. Error bars denote standard deviation (n = 3).

**Figure S16. Limiting glucose uptake capacity by pretreatment with GLUT1 inhibitor Bay876 sensitizes ATP levels to rotenone treatment in PC9 cells.** Cells were treated for 24h with either DMSO (control) or 20 uM Bay876, and then additionally with 1 uM Rotenone for 30 min. Data shown are ATP levels (as determined by CellTiter-Glo) relative to conditions without Rotenone treatment. Error bars denote standard deviation (n = 3). Experiments were performed in RPMI 1640 media with 5% FBS.

**Figure S17. ATP levels/cell upon acute treatment with metabolic effectors in PC9 cells.** ATP levels were determined with CellTiter-Glo (see methods), and normalized to untreated controls. Rotenone: complex I inhibitor. Oxamate: LDH inhibitor. Bay876: GLUT1 inhibitor. Oligomycin: ATP synthase inhibitor. FCCP: mitochondrial membrane uncoupler. Bromopyruvate: GAPDH inhibitor. Error bars denote standard deviation (n = 2). Experiments were performed in RPMI 1640 media with 5% FBS.

**Figure S18. Impact of small molecule effectors of metabolism on PC9 cell motility. A)**. Distribution of average PC9 cell speeds in individual tracks after monitoring cells for 24h. Black: DMSO control. Red: 12h pre-treatment with respective effector. Shown are distributions of three separate replicates. Red dashed line: median of distribution across all replicates. Black dashed line: median of average cell speed in DMSO. **B)** Median average speed plotted against respective growth rate. Error bars denote standard deviation (n = 3). All experiments were performed in RPMI 1640 media with 5% FBS.

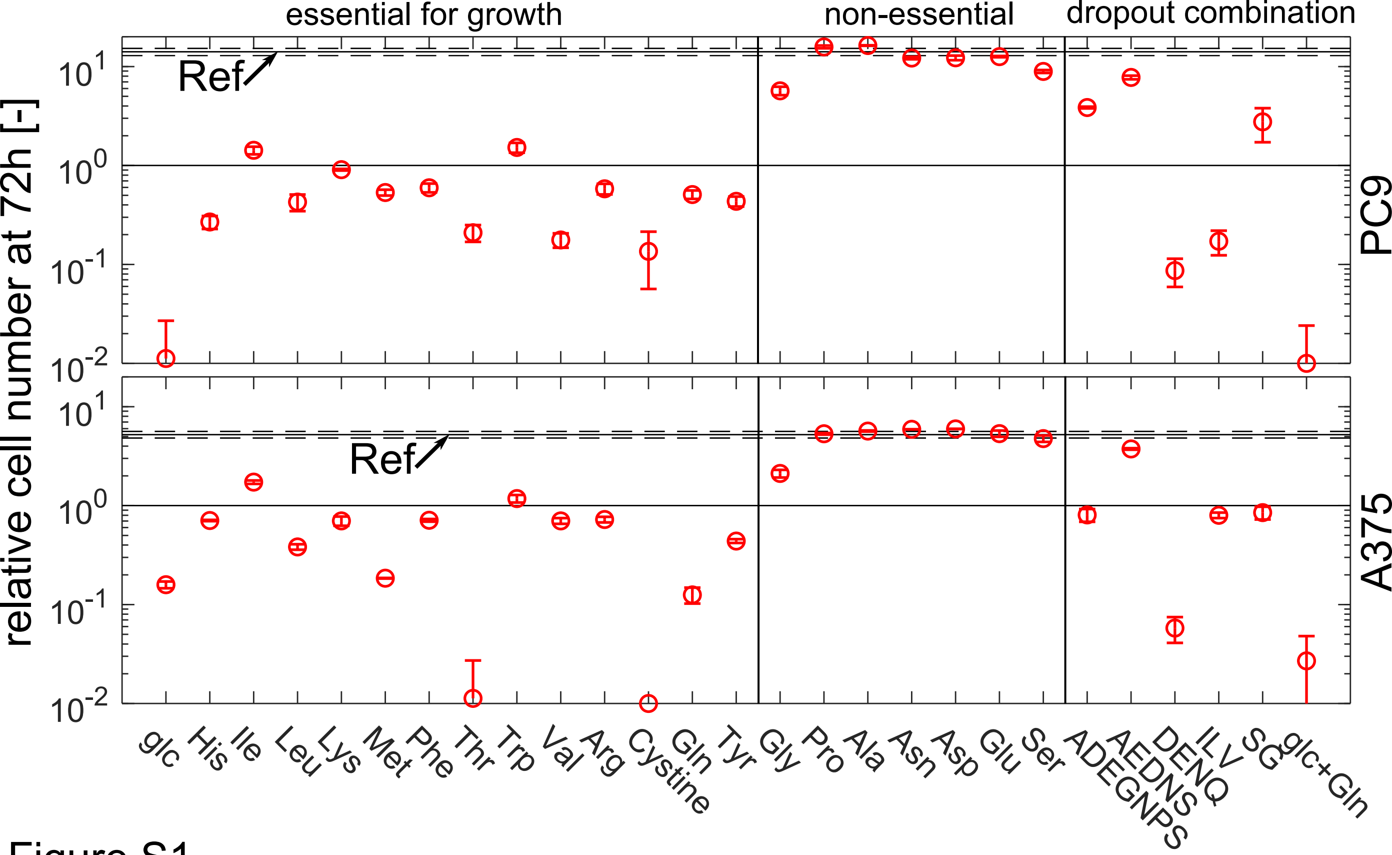
essential for growth
non-essential
dropout combination
relative cell number at 72h [-]
$10^{1}$
$10^{0}$
$10^{-1}$
$10^{-2}$
Ref
PC9
A375
glc
His
Ile
Leu
Lys
Met
Phe
Thr
Trp
Val
Arg
Cystine
Gln
Tyr
Gly
Pro
Ala
Asn
Asp
Glu
Ser
ADEGNPS
AEDNS
DENQ
ILV
SG
glc+Gln

Figure S1

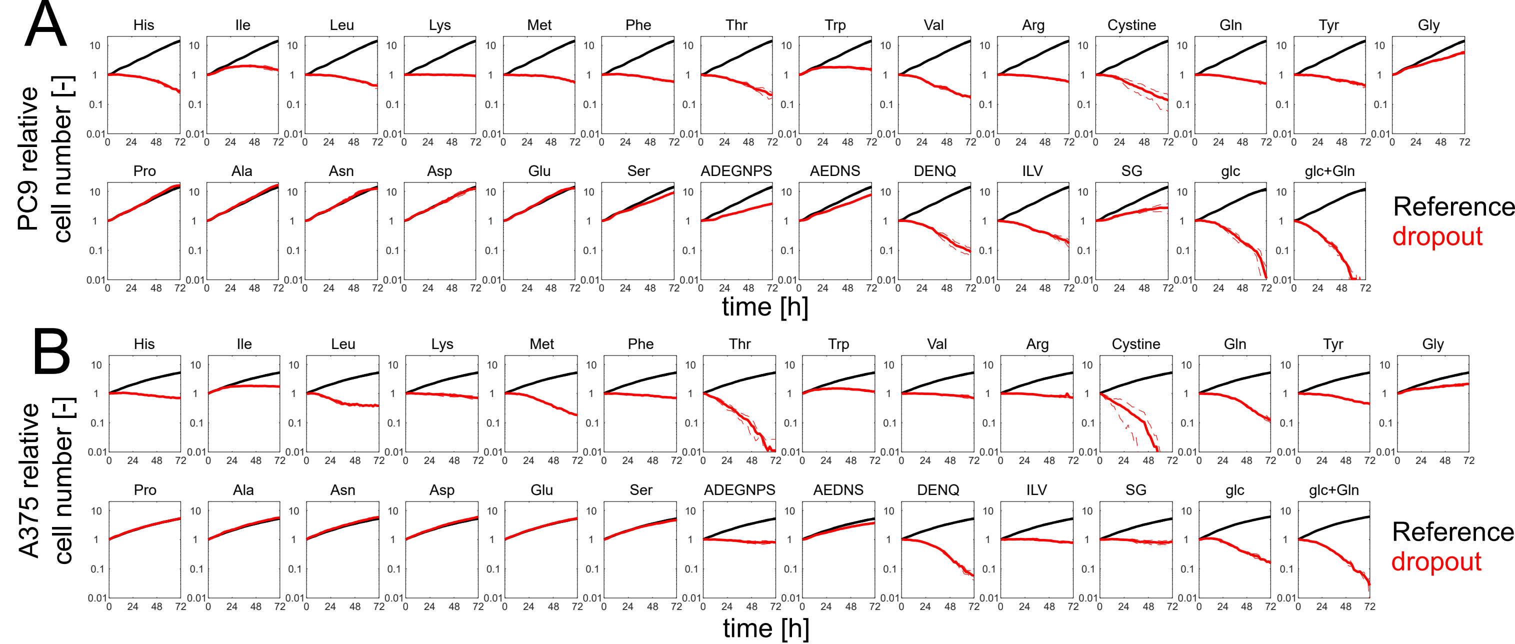
A
B
PC9 relative cell number [-]
A375 relative cell number [-]
His
Ile
Leu
Lys
Met
Phe
Thr
Trp
Val
Arg
Cystine
Gln
Tyr
Gly
Pro
Ala
Asn
Asp
Glu
Ser
ADEGNPS
AEDNS
DENQ
ILV
SG
glc
glc+Gln
10
1
0.1
0.01
0
24
48
72
time [h]
Reference
dropout

Figure S2

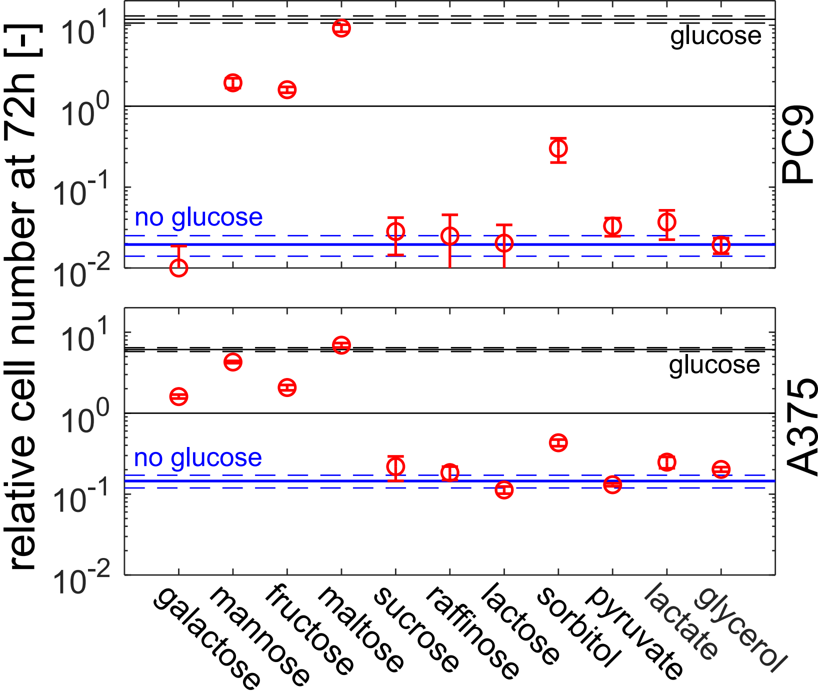
relative cell number at 72h [-]
$10^1$
$10^0$
$10^{-1}$
$10^{-2}$
glucose
no glucose
PC9
A375
galactose
mannose
fructose
maltose
sucrose
raffinose
lactose
sorbitol
pyruvate
lactate
glycerol

Figure S3

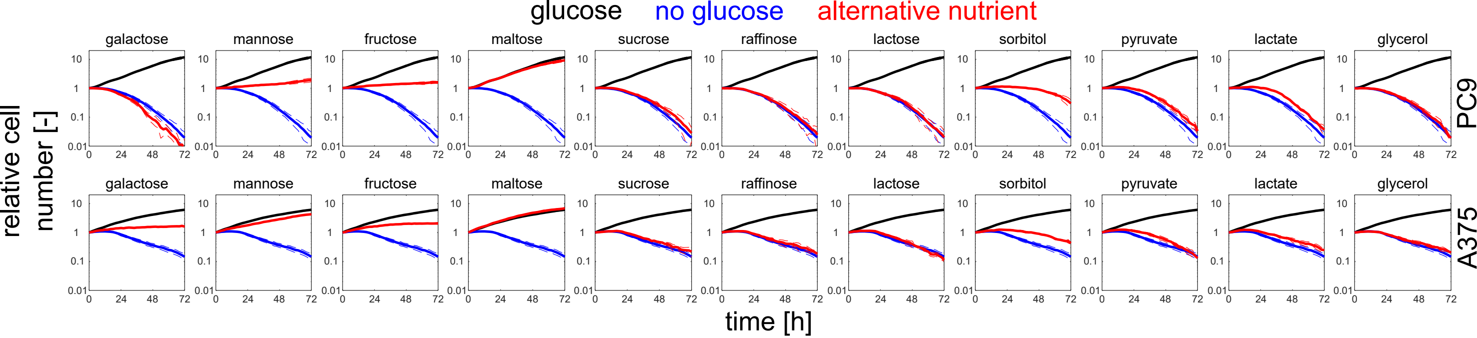
glucose
no glucose
alternative nutrient
galactose
mannose
fructose
maltose
sucrose
raffinose
lactose
sorbitol
pyruvate
lactate
glycerol
PC9
A375
relative cell number [-]
time [h]
0.01
0.1
1
10
0
24
48
72

Figure S4

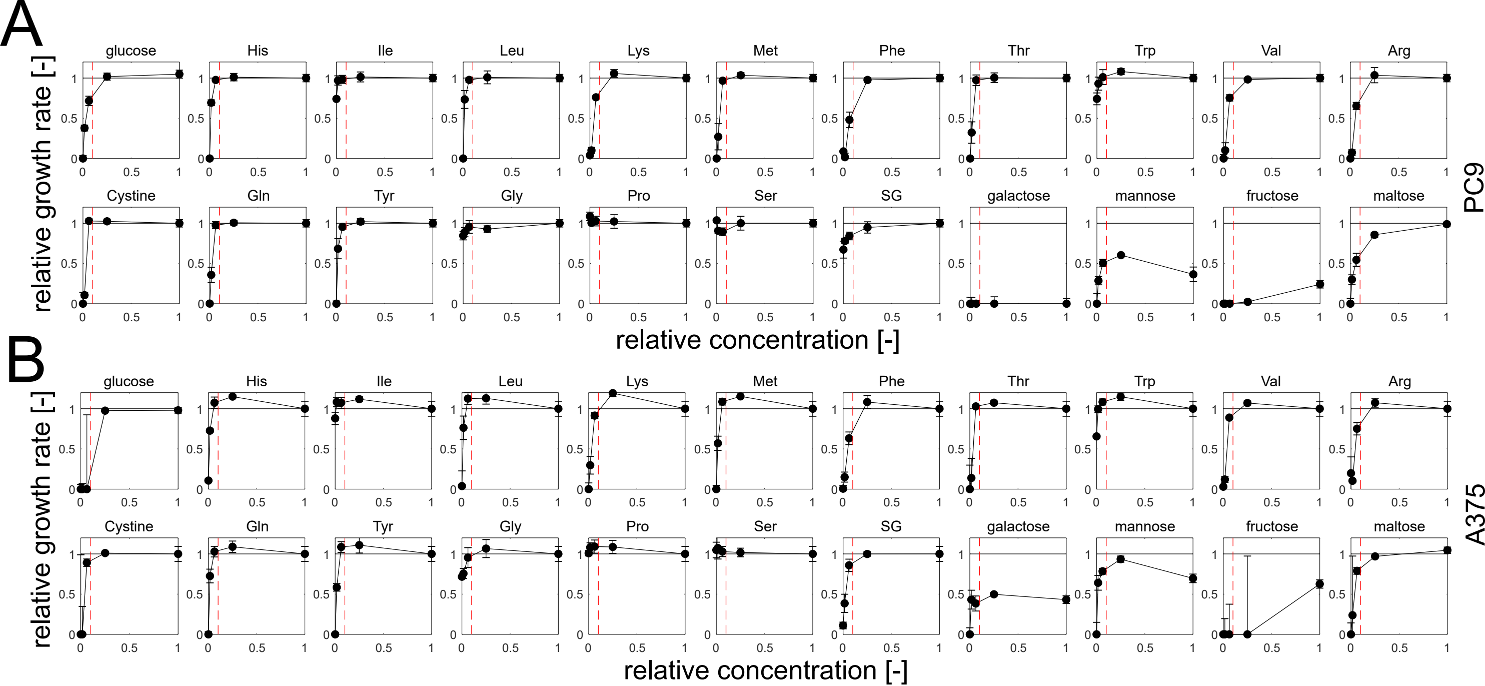

A
B
glucose
His
Ile
Leu
Lys
Met
Phe
Thr
Trp
Val
Arg
Cystine
Gln
Tyr
Gly
Pro
Ser
SG
galactose
mannose
fructose
maltose
relative growth rate [-]
relative concentration [-]
PC9
A375


Figure S5

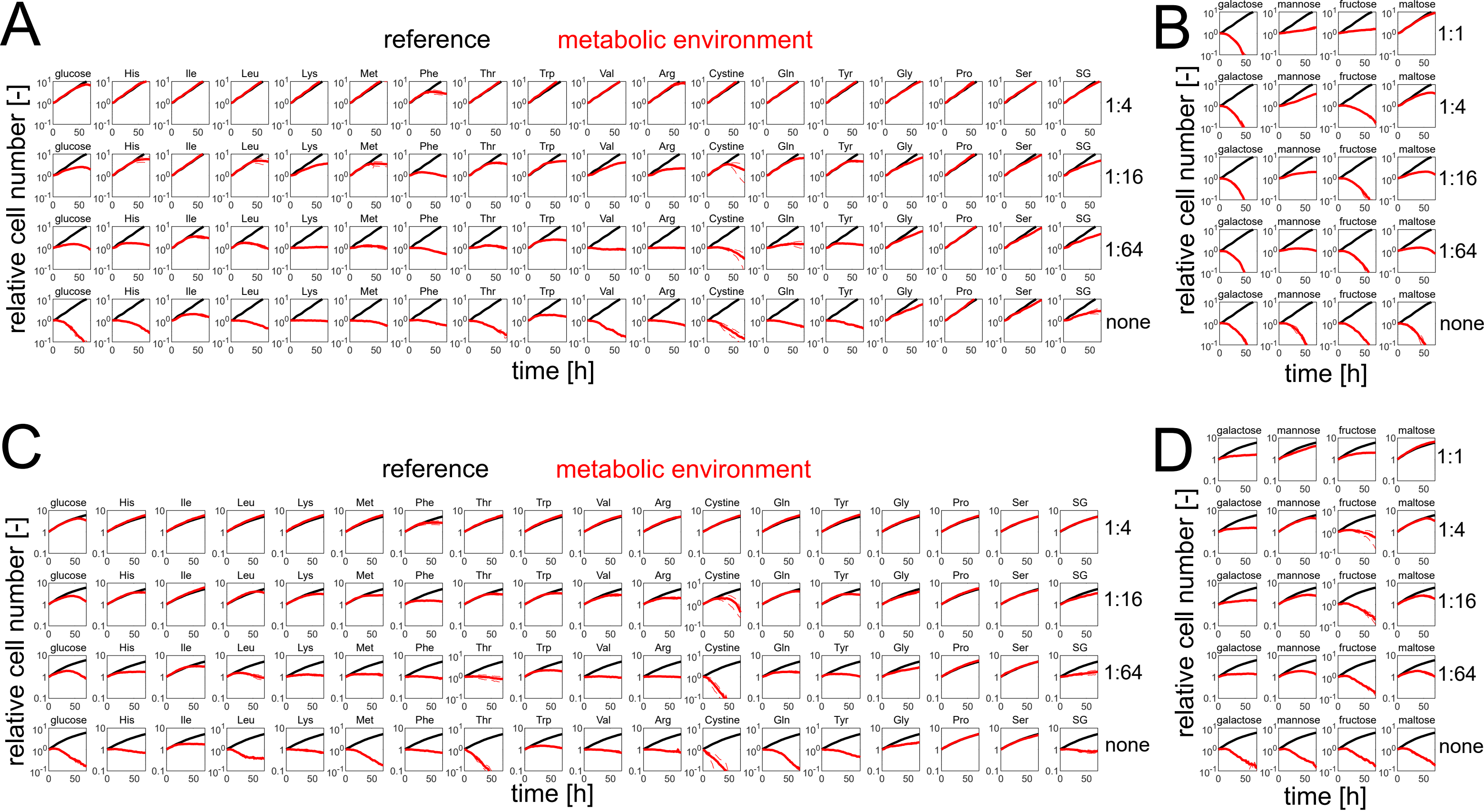
A
reference
metabolic environment
glucose
His
Ile
Leu
Lys
Met
Phe
Thr
Trp
Val
Arg
Cystine
Gln
Tyr
Gly
Pro
Ser
SG
1:4
1:16
1:64
none
relative cell number [-]
time [h]
B
galactose
mannose
fructose
maltose
1:1
1:4
1:16
1:64
none
relative cell number [-]
time [h]
C
reference
metabolic environment
glucose
His
Ile
Leu
Lys
Met
Phe
Thr
Trp
Val
Arg
Cystine
Gln
Tyr
Gly
Pro
Ser
SG
1:4
1:16
1:64
none
relative cell number [-]
time [h]
D
galactose
mannose
fructose
maltose
1:1
1:4
1:16
1:64
none
relative cell number [-]
time [h]

Figure S6

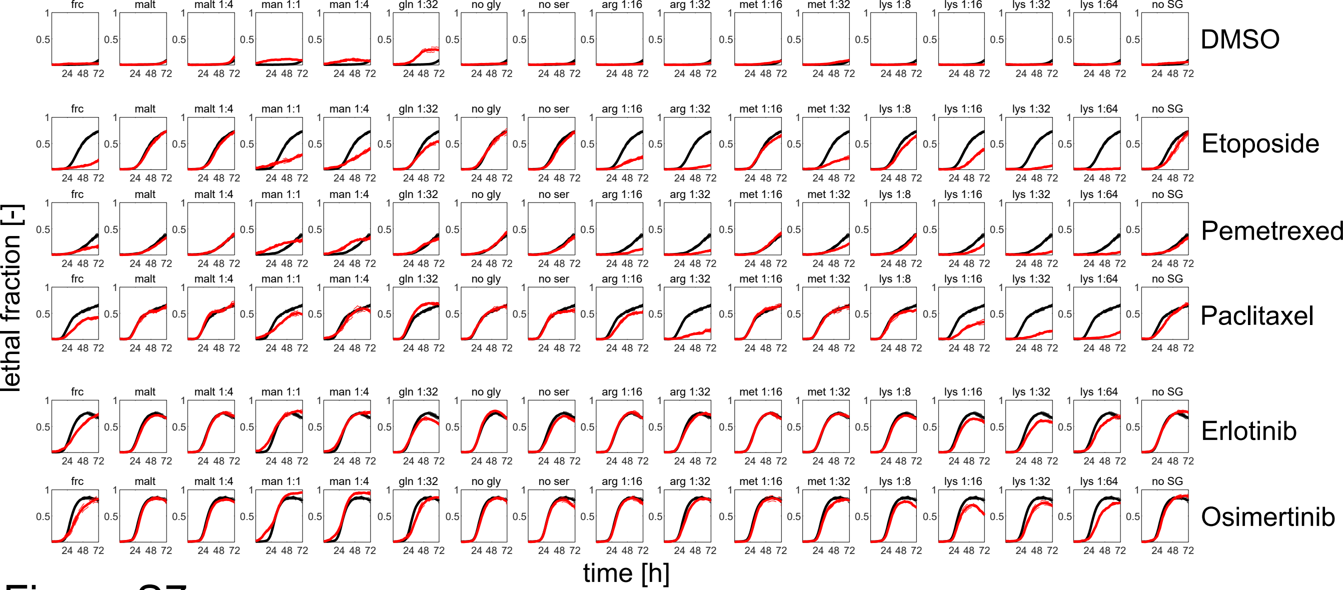
frc
malt
malt 1:4
man 1:1
man 1:4
gln 1:32
no gly
no ser
arg 1:16
arg 1:32
met 1:16
met 1:32
lys 1:8
lys 1:16
lys 1:32
lys 1:64
no SG
DMSO
Etoposide
Pemetrexed
Paclitaxel
Erlotinib
Osimertinib
lethal fraction [-]
time [h]

Figure S7

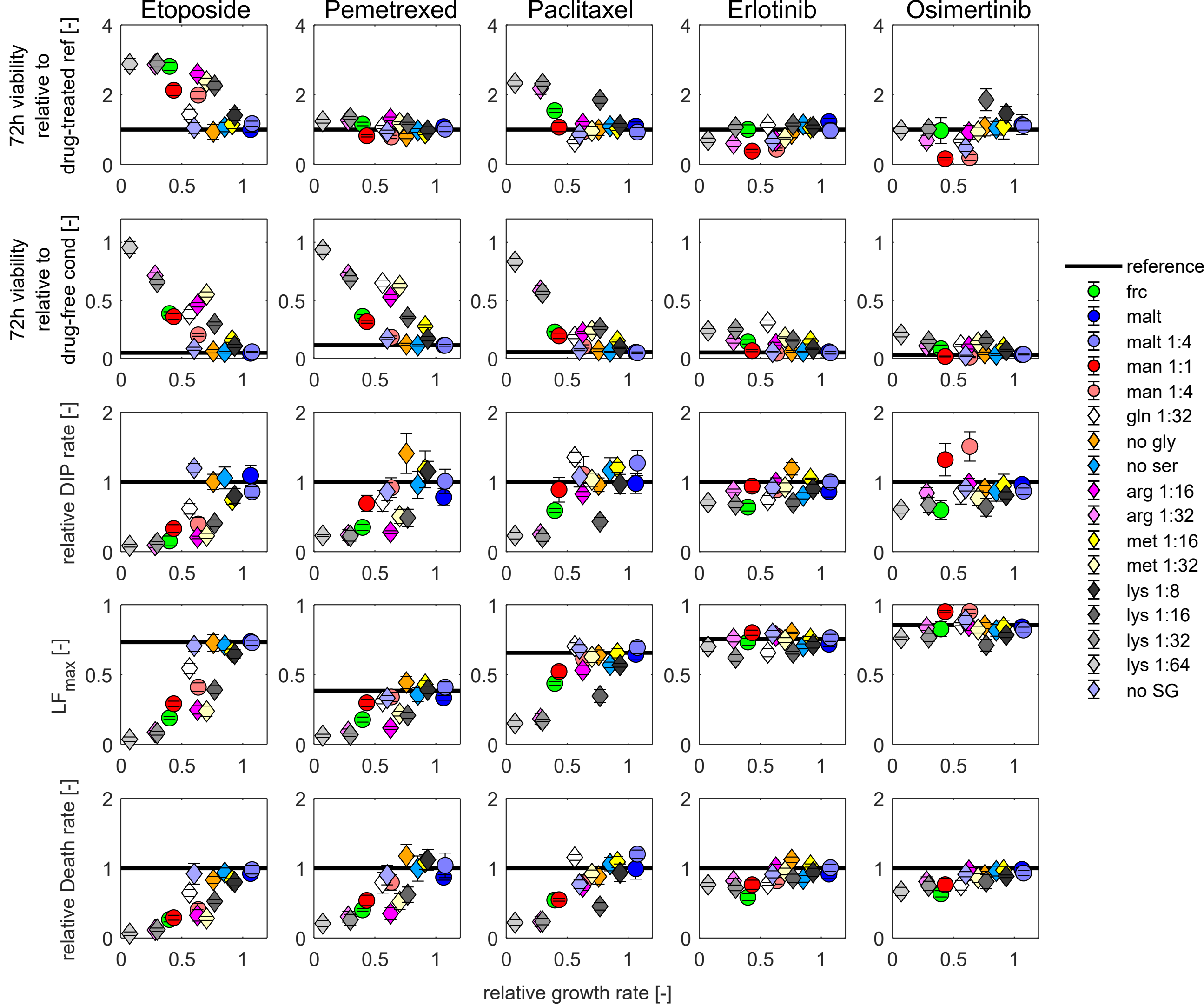
Etoposide
Pemetrexed
Paclitaxel
Erlotinib
Osimertinib
72h viability relative to drug-treated ref [-]
72h viability relative to drug-free cond [-]
relative DIP rate [-]
LF$_{max}$ [-]
relative Death rate [-]
relative growth rate [-]
reference
frc
malt
malt 1:4
man 1:1
man 1:4
gln 1:32
no gly
no ser
arg 1:16
arg 1:32
met 1:16
met 1:32
lys 1:8
lys 1:16
lys 1:32
lys 1:64
no SG

Figure S8

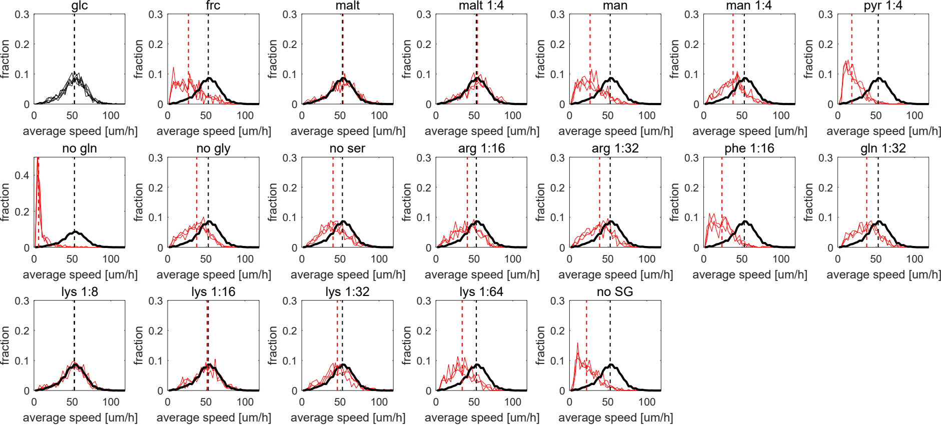
glc
frc
malt
malt 1:4
man
man 1:4
pyr 1:4
no gln
no gly
no ser
arg 1:16
arg 1:32
phe 1:16
gln 1:32
lys 1:8
lys 1:16
lys 1:32
lys 1:64
no SG
fraction
average speed [um/h]

### Figure S9

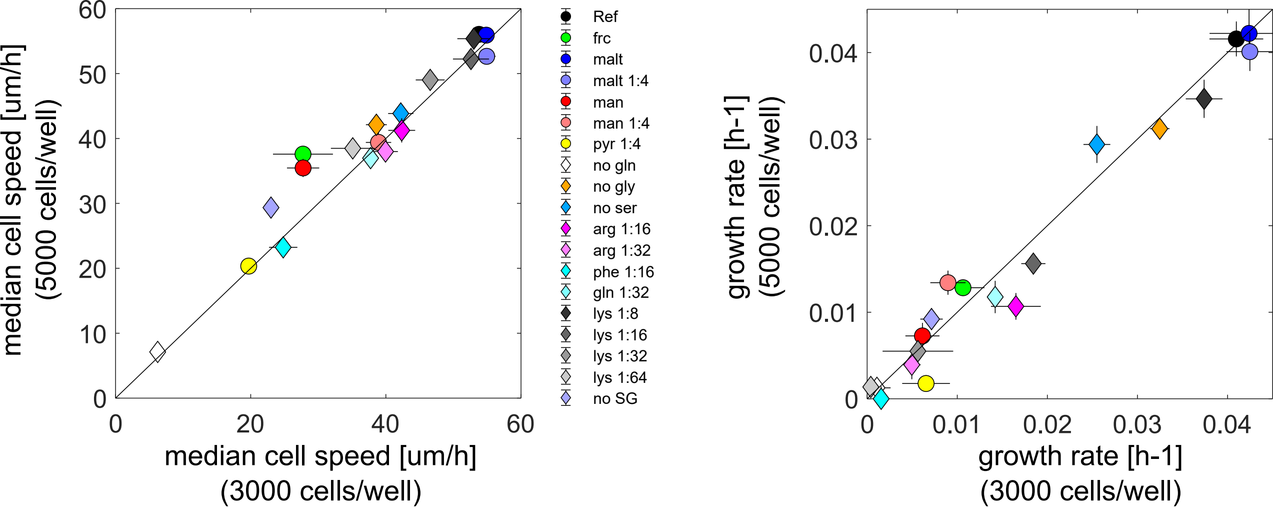

median cell speed [um/h] (5000 cells/well)
median cell speed [um/h] (3000 cells/well)
0
10
20
30
40
50
60
Ref
frc
malt
malt 1:4
man
man 1:4
pyr 1:4
no gln
no gly
no ser
arg 1:16
arg 1:32
phe 1:16
gln 1:32
lys 1:8
lys 1:16
lys 1:32
lys 1:64
no SG
growth rate [h-1] (5000 cells/well)
growth rate [h-1] (3000 cells/well)
0
0.01
0.02
0.03
0.04


Figure S10

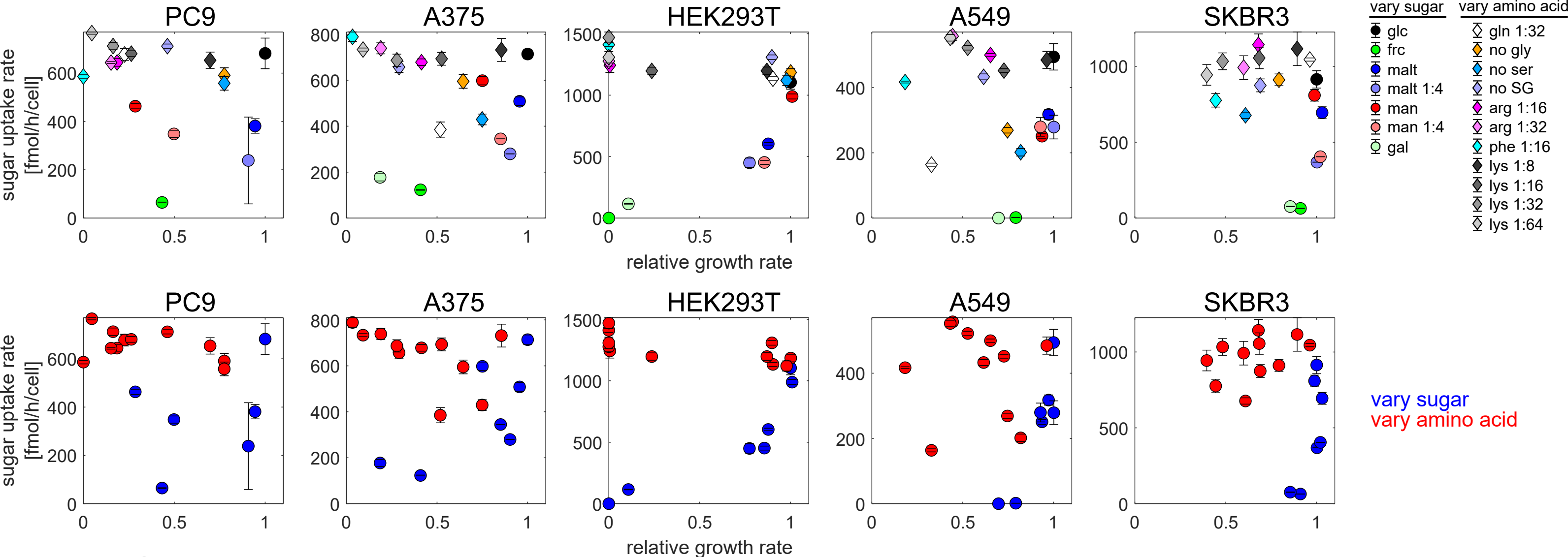
PC9
A375
HEK293T
A549
SKBR3
sugar uptake rate [fmol/h/cell]
relative growth rate
vary sugar
glc
frc
malt
malt 1:4
man
man 1:4
gal
vary amino acid
gln 1:32
no gly
no ser
no SG
arg 1:16
arg 1:32
phe 1:16
lys 1:8
lys 1:16
lys 1:32
lys 1:64
vary sugar
vary amino acid

Figure S11

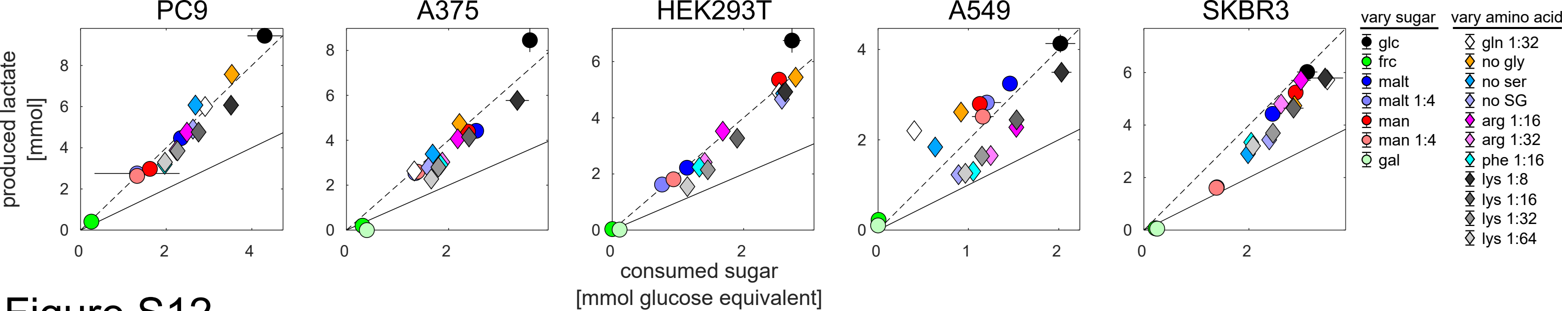

PC9
A375
HEK293T
A549
SKBR3
produced lactate
[mmol]
consumed sugar
[mmol glucose equivalent]
vary sugar
glc
frc
malt
malt 1:4
man
man 1:4
gal
vary amino acid
gln 1:32
no gly
no ser
no SG
arg 1:16
arg 1:32
phe 1:16
lys 1:8
lys 1:16
lys 1:32
lys 1:64


### Figure S12

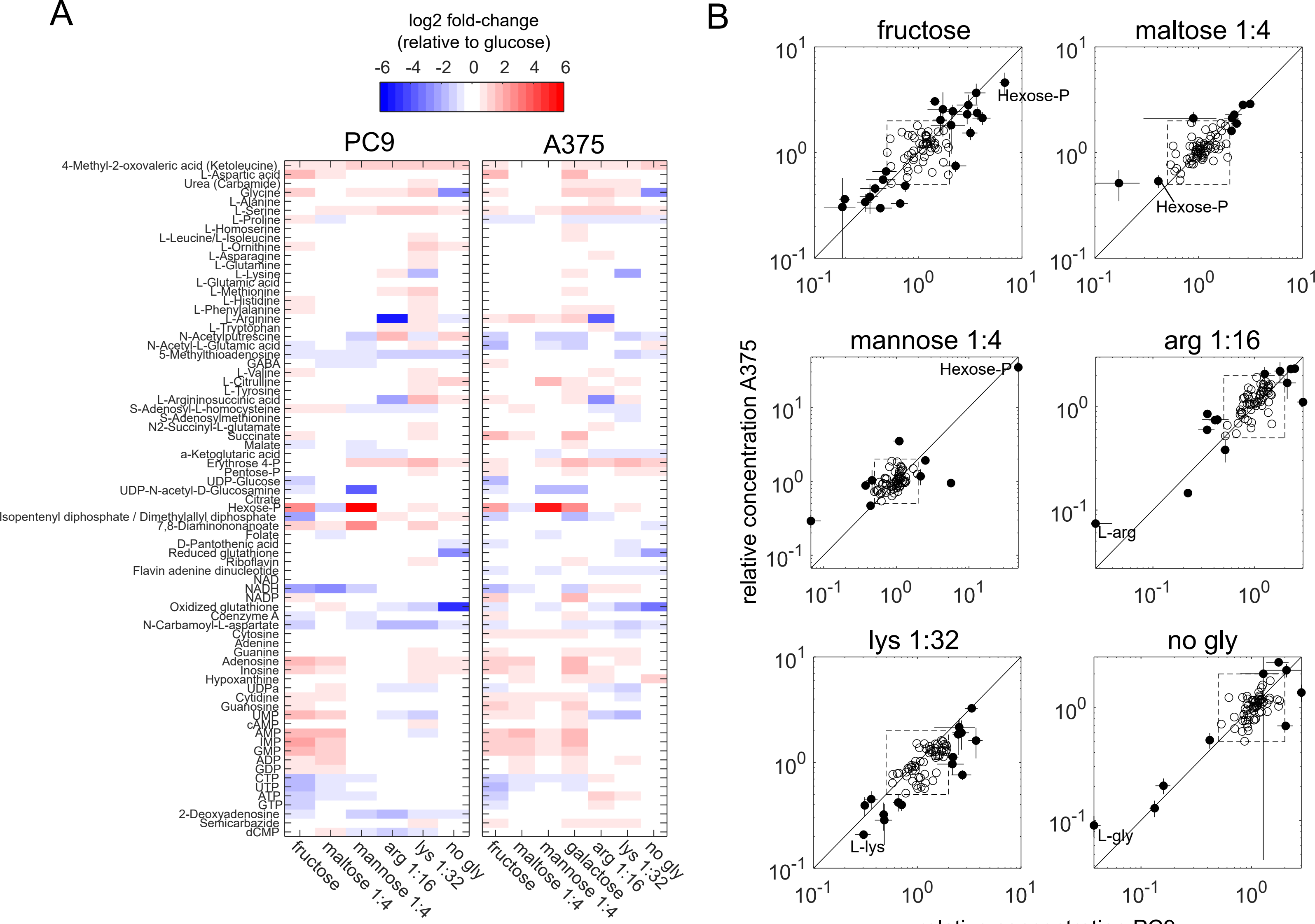

A
log2 fold-change
(relative to glucose)
-6 -4 -2 0 2 4 6
PC9
A375
fructose
maltose 1:4
mannose 1:4
galactose 1:4
arg 1:16
lys 1:32
no gly
B
fructose
maltose 1:4
mannose 1:4
arg 1:16
lys 1:32
no gly
Hexose-P
L-arg
L-lys
L-gly
relative concentration A375
relative concentration PC9


Figure S13

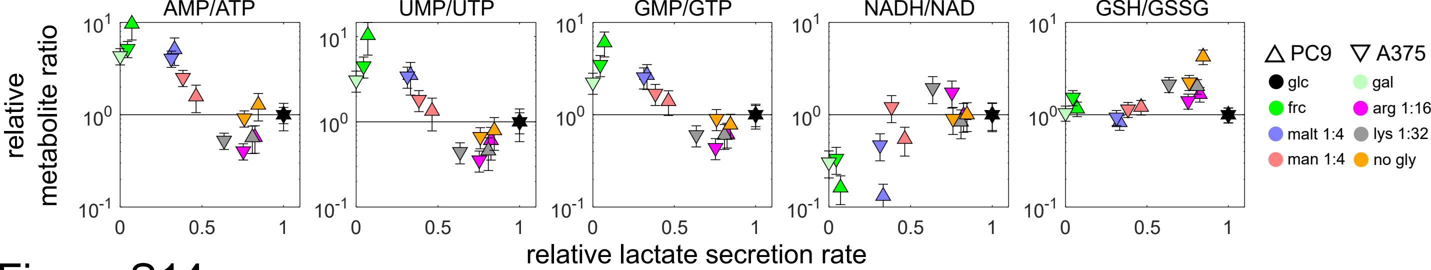

AMP/ATP
UMP/UTP
GMP/GTP
NADH/NAD
GSH/GSSG
relative metabolite ratio
relative lactate secretion rate
PC9
A375
glc
gal
frc
arg 1:16
malt 1:4
lys 1:32
man 1:4
no gly


Figure S14

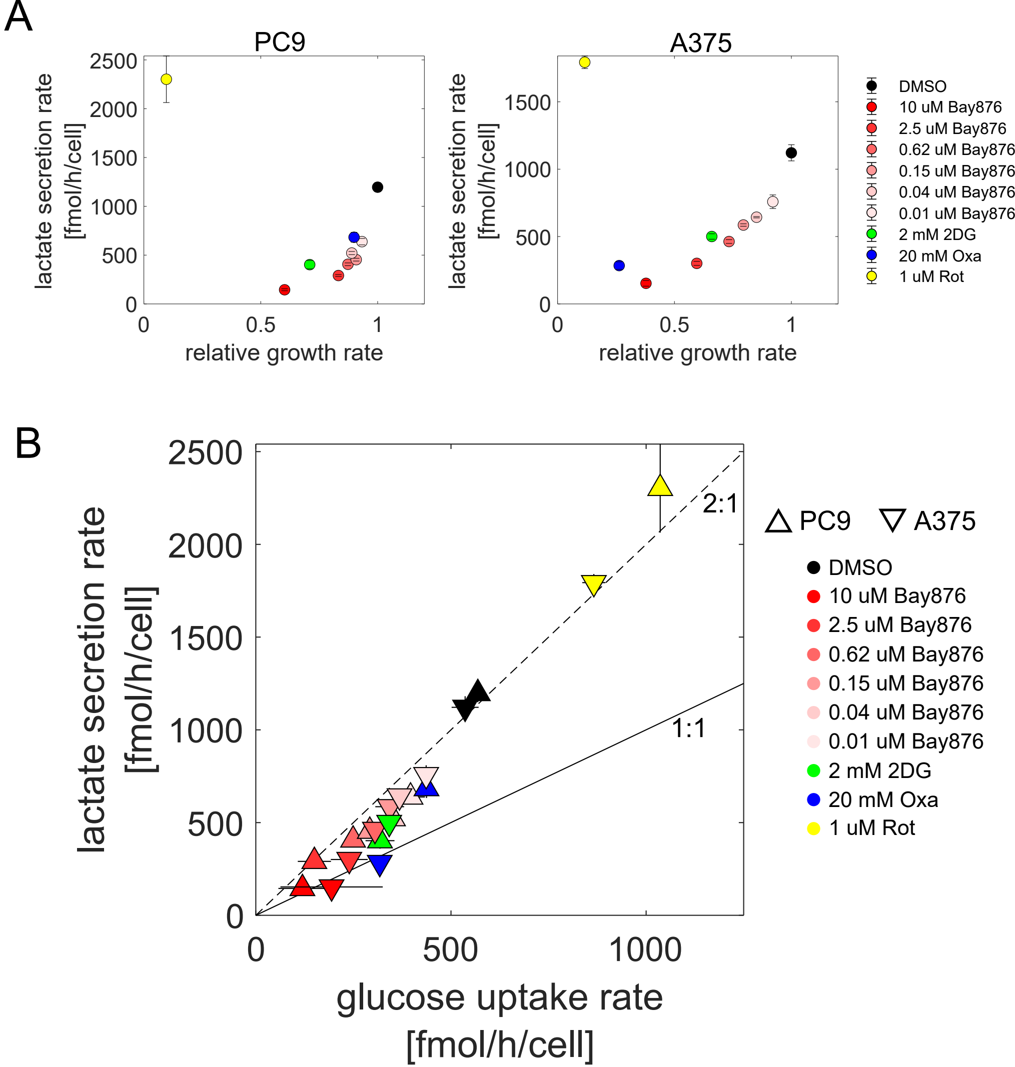
A
PC9
A375
lactate secretion rate [fmol/h/cell]
relative growth rate
DMSO
10 uM Bay876
2.5 uM Bay876
0.62 uM Bay876
0.15 uM Bay876
0.04 uM Bay876
0.01 uM Bay876
2 mM 2DG
20 mM Oxa
1 uM Rot
B
2:1
1:1
PC9
A375
glucose uptake rate [fmol/h/cell]

Figure S15

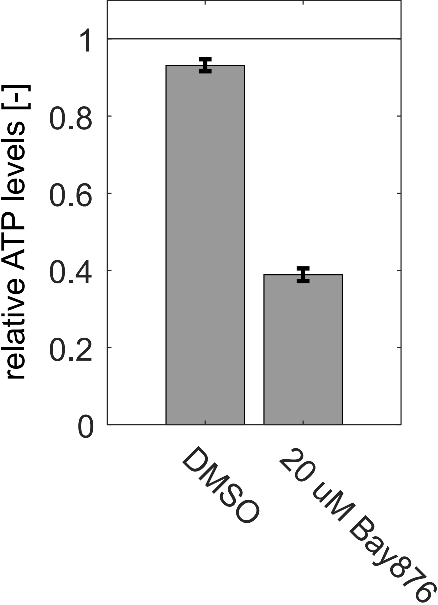


Figure S16

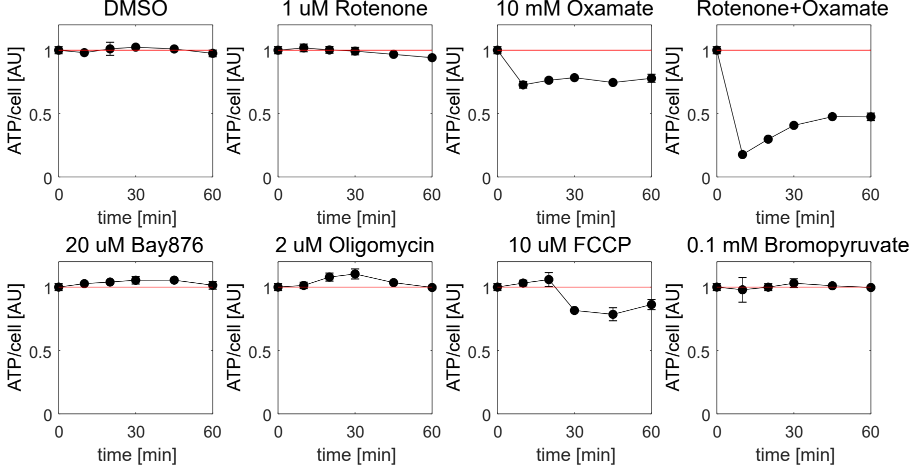
DMSO
1 uM Rotenone
10 mM Oxamate
Rotenone+Oxamate
20 uM Bay876
2 uM Oligomycin
10 uM FCCP
0.1 mM Bromopyruvate
ATP/cell [AU]
time [min]
0
0.5
1
0
30
60

### Figure S17

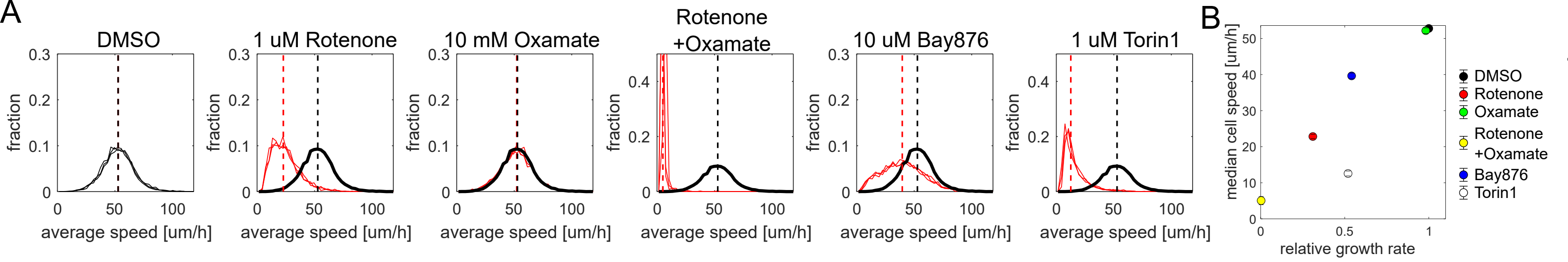
A
DMSO
1 uM Rotenone
10 mM Oxamate
Rotenone
+Oxamate
10 uM Bay876
1 uM Torin1
fraction
average speed [um/h]
B
median cell speed [um/h]
relative growth rate
DMSO
Rotenone
Oxamate
Rotenone
+Oxamate
Bay876
Torin1

Figure S18
